# Genome Wide Variant Analysis of Simplex Autism Families with an Integrative Clinical-Bioinformatics Pipeline

**DOI:** 10.1101/019208

**Authors:** Laura T. Jiménez-Barrón, Jason A. O’Rawe, Yiyang Wu, Margaret Yoon, Han Fang, Ivan Iossifov, Gholson J. Lyon

## Abstract

Autism spectrum disorders (ASD) are a group of developmental disabilities that affect social interaction, communication and are characterized by repetitive behaviors. There is now a large body of evidence that suggests a complex role of genetics in ASD, in which many different loci are involved. Although many current population scale genomic studies have been demonstrably fruitful, these studies generally focus on analyzing a limited part of the genome or use a limited set of bioinformatics tools. These limitations preclude the analysis of genome-wide perturbations that may contribute to the development and severity of ASD-related phenotypes. To overcome these limitations, we have developed and utilized an integrative clinical and bioinformatics pipeline for generating a more complete and reliable set of genomic variants for downstream analyses. Our study focuses on the analysis of three simplex autism families consisting of one affected child, unaffected parents, and one unaffected sibling. All members were clinically evaluated and widely phenotyped. Genotyping arrays and whole genome sequencing were performed on each member, and the resulting sequencing data were analyzed using a variety of available bioinformatics tools. We searched for rare variants of putative functional impact that were found to be segregating according to de-novo, autosomal recessive, x-linked, mitochondrial and compound heterozygote transmission models. The resulting candidate variants included three small heterozygous CNVs, a rare heterozygous *de novo* nonsense mutation in *MYBBP1A* located within exon 1, and a novel *de novo* missense variant in *LAMB3*. Our work demonstrates how more comprehensive analyses that include rich clinical data and whole genome sequencing data can generate reliable results for use in downstream investigations. We are moving to implement our framework for the analysis and study of larger cohorts of families, where statistical rigor can accompany genetic findings.

## Introduction

In 2010, the Center for Disease Control and Prevention (CDC) found that 1 in 68 eight-year-olds were diagnosed with ASD in the United States across 11 surveyed locations [1], with males being diagnosed 5 times more often than females [2-4]. Although the prevalence of ASD across different ethnicities, countries and social groups appears to be heavily influenced by methodological variables during diagnosis [5], it is clear that ASD is an emerging public health concern. Studies contributing to a better understanding of its causes and mechanisms promise to enable more precise diagnoses [6, 7], more effective treatments, and preventative care.

There is a vast and consistent amount of evidence suggesting a complex role of genetics in ASD [2, 6-12], in which many different loci are involved, but a general understanding of what causes ASD on a molecular and physiological level has not yet emerged. This question is broadly studied [2, 13-15], but the diversity of approaches used towards answering it has not led to broad conclusions about its etiology. Indeed, there is a large collection of putative disease contributing variants found in ASD diagnosed people, yet only a small fraction of these variants are reliably detected in small subpopulations of ASD patients [6, 7, 16, 17], leaving most ASD cases of undetermined etiology. The lack of generality in these findings may be attributed to many factors, including the phenotypic heterogeneity of the disease [8], the need for larger sample sizes for statistical studies [13], and to variability in the methodology used to analyze ASD-related data.

Currently, many ASD studies focus on the analysis of microarray and/or exome sequencing data for understanding the etiological contributions to and mechanisms of ASD [4, 9, 13]. These analyses are generally applied to large cohorts, such as those from the Simons Simplex Collection [4, 18], which consists of families with a single affected child, unaffected parents and at least one unaffected sibling. Those studies generally use and analyze only one of the high throughput sequencing technologies mentioned above, with varying levels of sequence coverage for WES or genotyping markers (for genotyping microarrays). Furthermore, these studies use only one or a few analysis tools for detecting sequence variations, which can result in a loss of information in situations where one tool performs poorly. Although these approaches have led to significant genetic discovery [4, 13], they are likely to miss-call or simply miss true and disease-relevant genetic variation. Some tools may perform better on just one or a few areas of the genome, and their performance may also differ depending on dataset-specific characteristics. To address this problem, we describe an integrative clinical and bioinformatics pipeline that makes use of a variety of analysis tools and orthogonal high throughput sequencing technologies to obtain a more complete and reliable set of candidate ASD-variants for validation and downstream functional analysis.

## Results

This study consisted of the clinical recruitment of three Simplex Autism Families (Figure 1) for phenotyping and whole genome studies. Human sequence variation spans a variety of genomic scales, ranging from single nucleotide to megabase and even whole chromosome differences. Due to the variety of scales and mechanisms that can lead to variation in human sequence between individuals and populations, a variety of algorithms are needed in order to extract genomic signatures at all scales and that represent a wide variety of variant types. A variety of variant discovery algorithms and procedures were used during the course of this study, each designed to detect different classes of human sequence variants (Figure 2).

**Figure 1.**
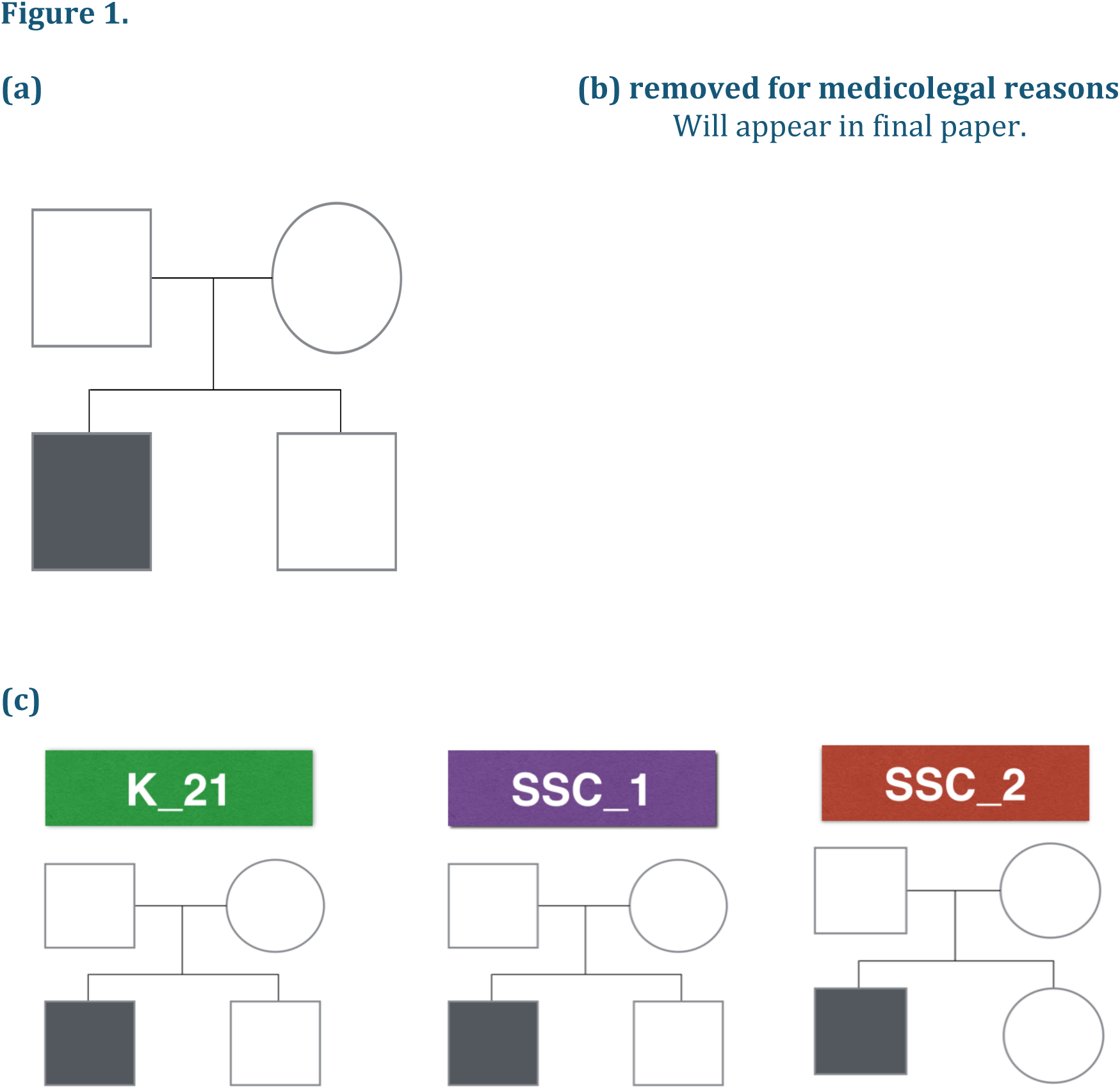
(a) Pedigree structure of a Simplex Autism Family. For a family to be classified as a Simplex Autism Family, it has to be composed of one affected child and at least one unaffected sibling, and both parents do not have obvious autism. Probands and siblings can be either males or females. **(b) K_21 Proband showing no dysmorphology. (c) Analyzed Pedigrees.** Two of the families have male probands and unaffected male siblings (K_21 and SSC_1), whereas the third family has a male proband and a female unaffected sibling (SSC_2).

**Figure 2.**
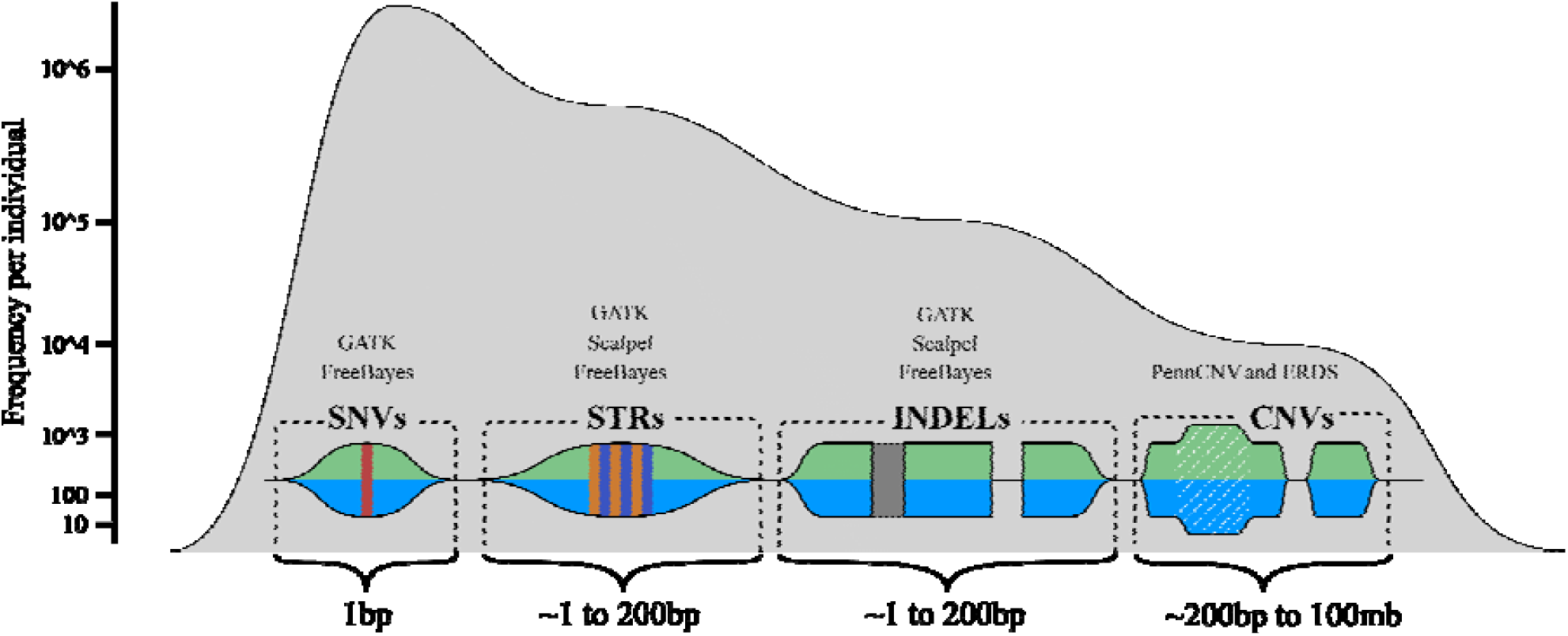
A conceptual map of human sequence variation. Here, we show approximate sizes, as well as the associated signature, of the various different types of human sequence variation that can be currently detected with the WGS, microarray sequencing and informatics technologies employed in this work. The frequency axis shows the approximate frequency of the various genetic variation types that are currently detectable via germline WGS combined with microarray sequencing. Above the visual signatures of the different types of human sequence variation, the general names of the different informatics software tools for detecting the variation are noted which include, the Genome Analysis Took Kit (GATK), Scalpel, PennCNV, the estimation by read depth with single-nucleotide variants (ERDS) CNV caller and the FreeBayes caller.

### Simon’s Simplex Collection Phenotypic evaluation results

Body Mass Index (BMI), head circumference, height and weight were measured for both probands stemming from the two Simons Simplex Collection (SSC) families studied here (Table 1). General IQ (intellectual quotient) as well as verbal IQ and non-verbal IQ were measured for both SSC probands. Table 2 details the evaluative instruments used for each SSC family. Similar body and cognitive measurements were not available for the third K22 proband (Table 1), although ancestral background was recorded for all three probands.

**Table 1.** Body measurements and IQ tests scores.

| Proband | BMI | Head Circumference | Height | Weight | Race | Proband IQ | Proband VIQ | Proband NVIQ |
| --- | --- | --- | --- | --- | --- | --- | --- | --- |
| SSC_2 | 17.4 | 51.6 | 102 | 18.1 | Asian | 61 | 32 | 89 |
| SSC_1 | 32.9 | 60.5 | 175 | 100.9 | Caucasian | 120 | 136 | 108 |
| K_21 | NA | NA | NA | NA | Caucasian | NA | NA | NA |

**Table 2.** The Aberrant Behavior Checklist scores for each Proband.

| Proband | ABC I: Irritability | ABC II: Lethargy | ABC III: Stereotypy | ABC IV: Hyperactivity | ABC V: Inappropriate Speech |
| --- | --- | --- | --- | --- | --- |
| SSC_2 | 8 | 13 | 5 | 27 | 0 |
| SSC_1 | 14 | 5 | 3 | 6 | 6 |
| K_21 | NA | NA | NA | NA | NA |

#### Concordance between variant detection algorithms

In this section, we explore detection reliability by measuring concordance among algorithm results across all sequenced individuals. Single nucleotide variants, small insertions or deletions and the detection of *de novo* variants of either class were compared across algorithms applied to raw whole genome sequencing (WGS) data.

### Single nucleotide variants (SNVs) and small insertions or deletions (INDELs)

GATK (the Genome Analysis Toolkit) and Freebayes are algorithms that detect both SNVs and INDELs across the entire sequenced genome; as such, we report here the concordance between these two algorithms in detecting SNVs and INDELs. The observed mean concordance between GATK and Freebayes was 79.3% and 56.6% for filtered SNV and INDEL calls, respectively. After filtering for high quality variants according to each algorithm’s recommendation (see methods), concordance between the algorithms increases by 5.7% and 5.4% for SNVs and INDELs respectively. Table 3 summarizes the mean per person number of variants called by each algorithm.

**Table 3.** The Autism Diagnostic Observation Schedule (ADOS) scores for each Simon’s Simplex Collection Proband.

| Proband | ADOS1 Algorithm | ADOS Communication | ADOS Reciprocal Social |
| --- | --- | --- | --- |
| SSC_2 | no-words | 7 | 11 |
| SSC_1 | NA | 3 | 8 |
| K_21 | NA | NA | NA |

### De-novo unique SNVs

The mean number of unfiltered unique *de novo* SNVs (not shared by siblings) detected by the Multinomial Analyzer, Freebayes and GATK was 65,572, 76,920 and 40,873 respectively. The Multinomial Analyzer is an algorithm specifically designed to detect *de novo* SNVs, where as additional steps were taken to obtain a list of putative *de novo* variants using Freebayes and GATK. After filtering variants based on each algorithm’s recommendations (refer to the Methods section for details pertaining to the filtering procedures), the mean number of variants detected by each caller dropped to 1,692, 24,982 and 31,831 for the Multinomial Analyzer, Freebayes and GATK, respectively. The concordance between the 3 algorithms was generally low, with Freebayes and GATK agreeing on 12.4% of their detected variants, and all three agreeing on less than one percent of the total filtered call-set, 0.113%. It is important to note that the low concordance between the Multinomial Analyzer and the other algorithms is influenced by the fact that its filtering step considers a ‘*de novo* score’, which is something that the other algorithms do not use for filtering purposes. Thus, the large difference in overall call rate makes a comparison of the mean overlapping calls somewhat uninformative, as the intersection between all three can only be as large as the smallest set. It is for this reason that the union of the three algorithms was considered during downstream prioritization steps, rather than the intersection.

### De-novo unique INDELs

*De-novo* INDELs from GATK, Freebayes and Scalpel were also compared. The mean number of *de novo* INDELs detected by GATK, Freebayes and Scalpel per proband before filtering was 52,631, 55,505 and 128 respectively, and after filtering based on each algorithm’s recommendations, this number dropped to 42,425, 37,210 and 70. The concordance between the three algorithms was, again, low. Freebayes and GATK agreed on 10.7% of the total call set, and all three callers agreed on only one variant and only within the subset of a single family (i.e., there was only one instance in which all three callers found the same variant). One should keep in mind that the filtering criteria and size of call sets are very different across these three callers, so our a-priori expectation is that a low number of calls will be within the intersection of all three.

#### Variant classification and prioritization for SNVs and INDELs

After obtaining high quality call sets from the union of filtered variants from all algorithms and categorizing them according to different disease models, the number of variants was still too large to proceed to more detailed literature searches and putative functional interpretations. Filtering variants by CADD score > 20 and MAF < 0.01 from 1000 genomes reduced the number of variants for consideration dramatically (Table 4) and the number of compound heterozygous mutations was reduced to zero. However, variants segregating according to the compound heterozygous model are not necessarily expected to be deleterious on their own, but may be deleterious in combination with other variants in the same gene on the same, or different, chromosome.

**Table 4.** The Child Behavior Checklist scores for each Simon’s Simplex Collection Proband.

| Proband | ADD/ ADHD | Aggressive Behavior | Anxiety Problems | Anxious/ Depressed | Attention Problems | Defiant | Withdrawn | Total Problems |
| --- | --- | --- | --- | --- | --- | --- | --- | --- |
| SSC_2 | 52 | 55 | 51 | 56 | 53 | 59 | 82 | 61 |
| SSC_1 | 57 | 70 | 58 | 63 | 51 | 66 | 63 | 66 |
| K_21 | NA | NA | NA | NA | NA | NA | NA | NA |

To narrow the list of possible disease contributing variants, each call set was annotated and filtered using various criteria and scores described in the Methods section. Out of the resulting prioritized variants, an average of 101 per family were localized to intra or intergenic regions (**Supp. Table 1**) and only three were located within a gene, one of which was found to be common in the SSC controls. Thus, by these filtering criteria, two exonic variants were considered as potentially contributing to the disease (Table 5). The genic variants are described below.

**Table 5.** The Vineland Adaptive Behavior Scale scores for each Simon’s Simplex Collection Proband.

| Proband | Communication | Daily Living Skills | Motor Skills | Socialization |
| --- | --- | --- | --- | --- |
| SSC_2 | 76 | 58 | 100 | 63 |
| SSC_1 | 81 | 89 | NA | 73 |
| K_21 | NA | NA | NA | NA |

### MYBBP1A stop gain variant

A *de novo* heterozygous nonsense mutation was found on the first exon of *MYBBP1A* (chr17: 4,442,191-4,458,926) in pedigree K21 (Figure 3). This mutation is located at chr17:4458481, it is a G->A substitution and is annotated as being highly deleterious with a CADD score of 40, which corresponds to being within the top 0.01% of all possible SNVs in terms of its deleteriousness. The variant was not found in DBsnp Human Build 142 [19], the exome variant server [http://evs.gs.washington.edu/EVS/] or in any other person in the Simon’s Simplex Collection database. One proband from the SSC was found to have a *de novo* missense G->T substitution in the same gene located at chr17: 4444853 causing an Arg->Ser change. Only one person out of 71,164 unrelated individuals from the Exome Aggregated Consortium [http://exac.broadinstitute.org] is reported to have this exact same mutation, indicating that this is a very rare variant. As the phenotype of this person in the ExAC database with the mutation is unknown, and also given that there are people with neuropsychiatric conditions in ExAC, no conclusions can be made from this alone. Sanger sequencing validated this mutation (Supp. Fig. 1).

**Figure 3.**
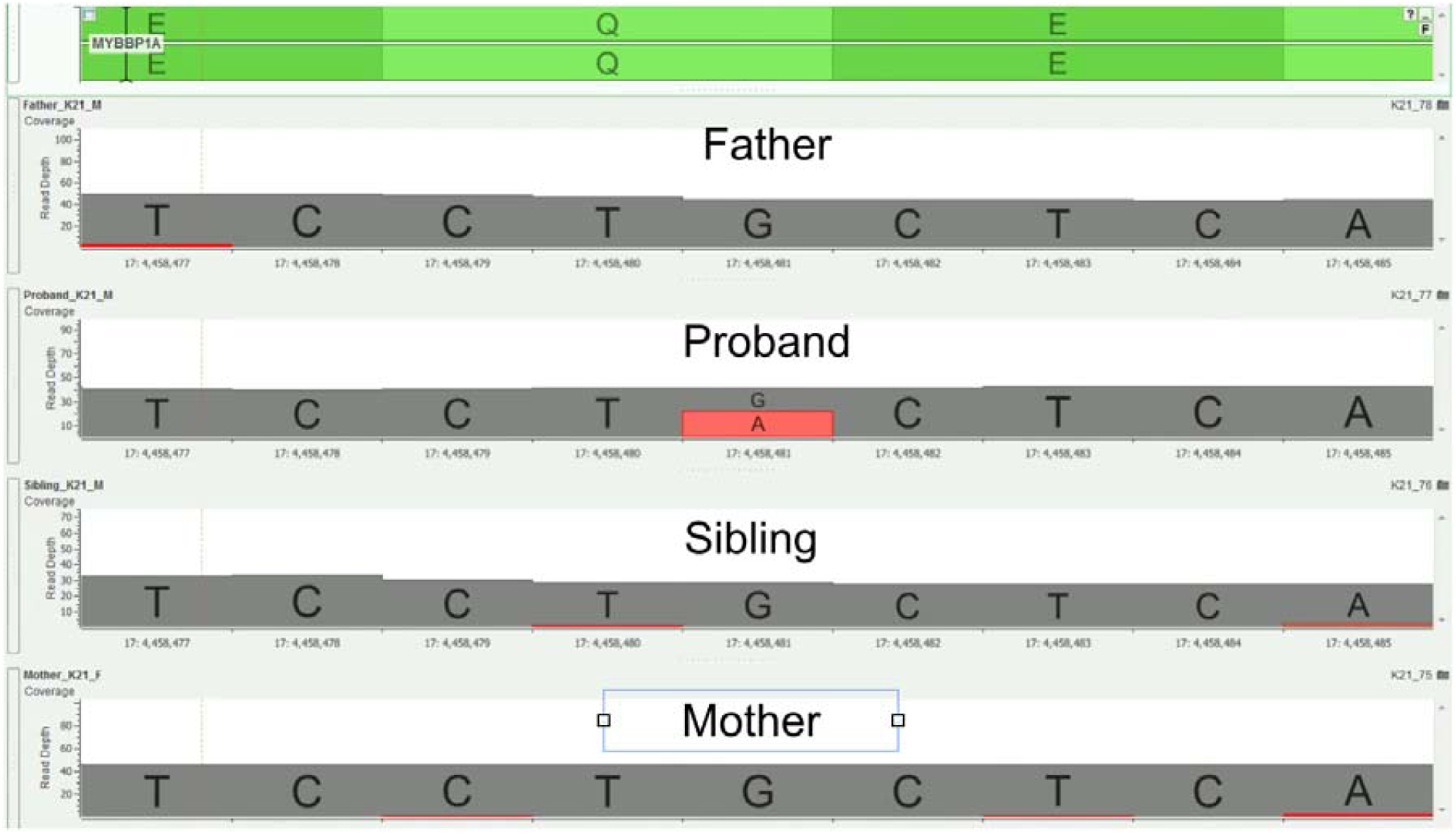
Genome Browser Screen cut for the read depths in the **MYBBP1A stop gain.** (chr17:4458481) mutation in K21 family.

### LAMB3 missense variant

The second *de novo* mutation detected in the study was found in the SSC_1 pedigree, this time a missense mutation located at chr1:209823359 on *LAMB3* (Figure 4). Although this mutation was reported in a previous study [13], it was not found in any other person contained within the SSC database and it was not found in any of the other interrogated databases, the exome variant server, or the ExAC database, making it an ultra rare mutation.

**Figure 4.**
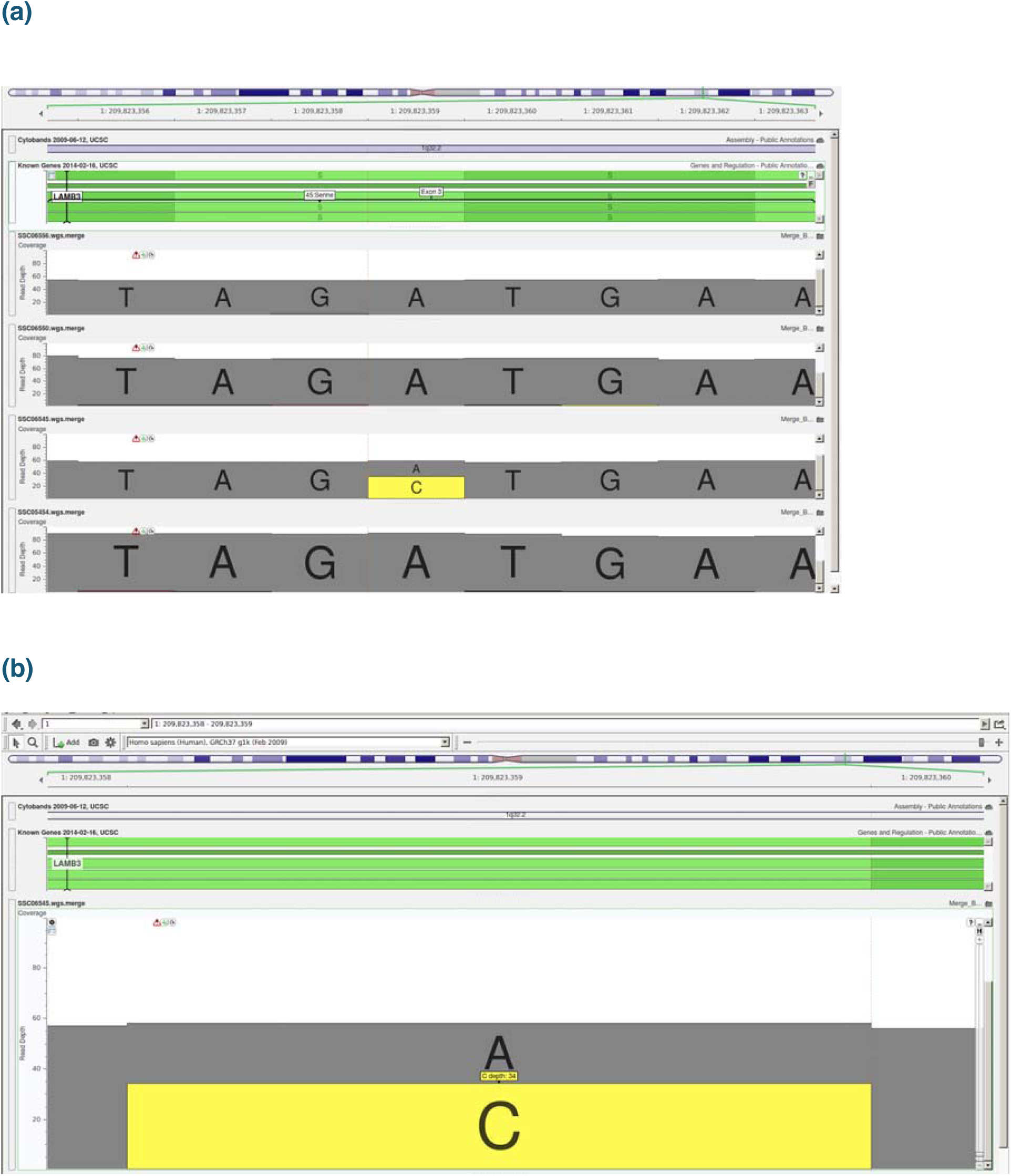
**(a).** Genome Browser Screen cut for the read depths in the LAMB3 missense mutation (chr1:209823359). **(b).** Genome Browser screen cut showing 34 reads supporting the variant for the proband in SSC_1 family.

#### Variant classification and prioritization for copy number variations

On average, 1500 unfiltered deletions and 450 unfiltered duplications were detected by ERDS applied to the WGS data (see Methods) for each person in the study. After filtering (see Methods), 150 deletions and 170 duplications were found on average per person. The number of calls obtained with PennCNV was highly variable, with a mean of 60 unfiltered duplications (sd=38) and a mean of 80 unfiltered deletions (sd=29) being detected. After filtering the variants, only 5% and 20% of all duplications deletions were retained, respectively. After annotating, none of the remaining CNVs were identified as pathogenic. However, we detected three CNVs (Figure 5-7) whose coordinates (Table 6) are embedded within larger CNVs that have been associated with cognitive disease. These CNVs were not found in any other unaffected family member. Two out of the three CNVs were found in pedigree K21, however only the ERDS algorithm detected them. As described in the Methods section, PennCNV uses the Log R Ratio (LRR) and B Allele Frequencies (BAF) to detect a CNV. Different numbers of copies have different clustering patterns for the LRR and BAF values when plotted. In pedigree K21 (Figure 5, 6), both the LRR and BAF are not properly clustered, suggesting, in this case, that these CNVs were not detected by PennCNV but were detected by ERDS as true positives, due to the properties of the microarray dataset for this family.

**Figure 5.**
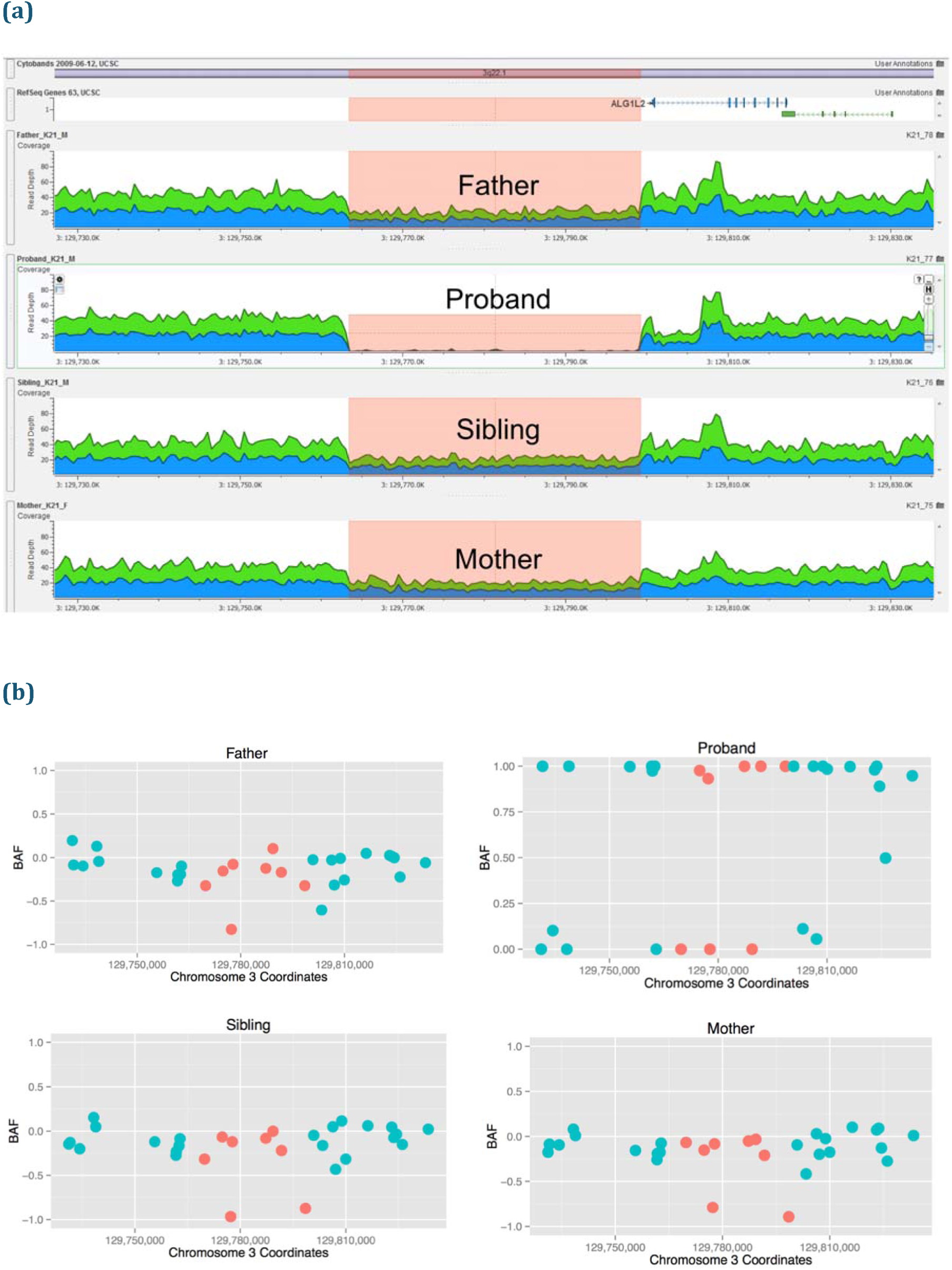
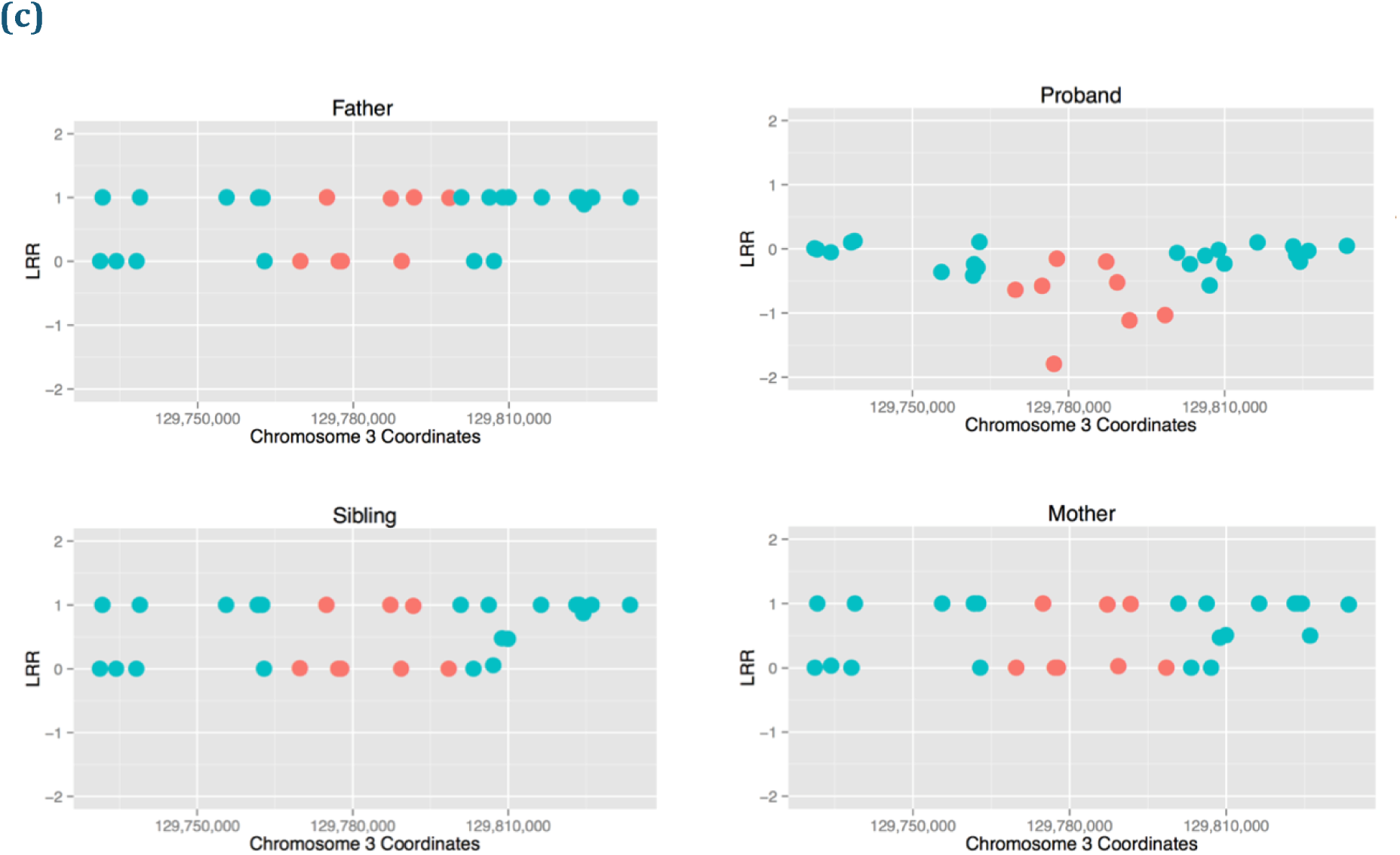
**(a).** Genome Browser Screen cut for the Read Depths in the K21 CNV 3q22.1 region of 16Kb. **(b).** B Allele Frequencies values for Illumina Omni 2.5 markers on 3q22.1 region including the markers belonging to the CNV region detected by ERDS in red. **(c).** Log R Ratio values for Illumina Omni 2.5 markers on 3q22.1 region including the markers belonging to the CNV region detected by ERDS in red.

**Figure 6.**
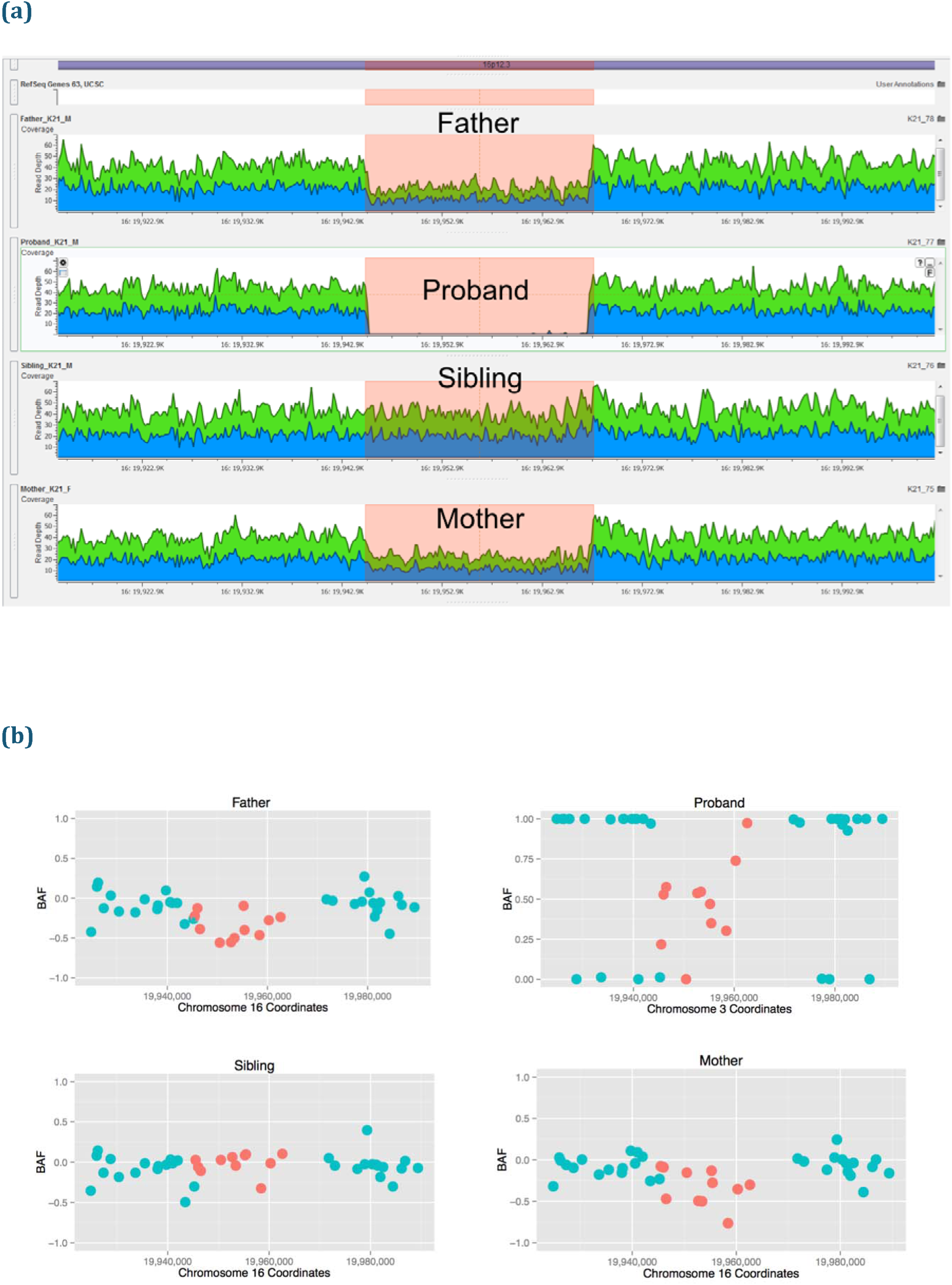
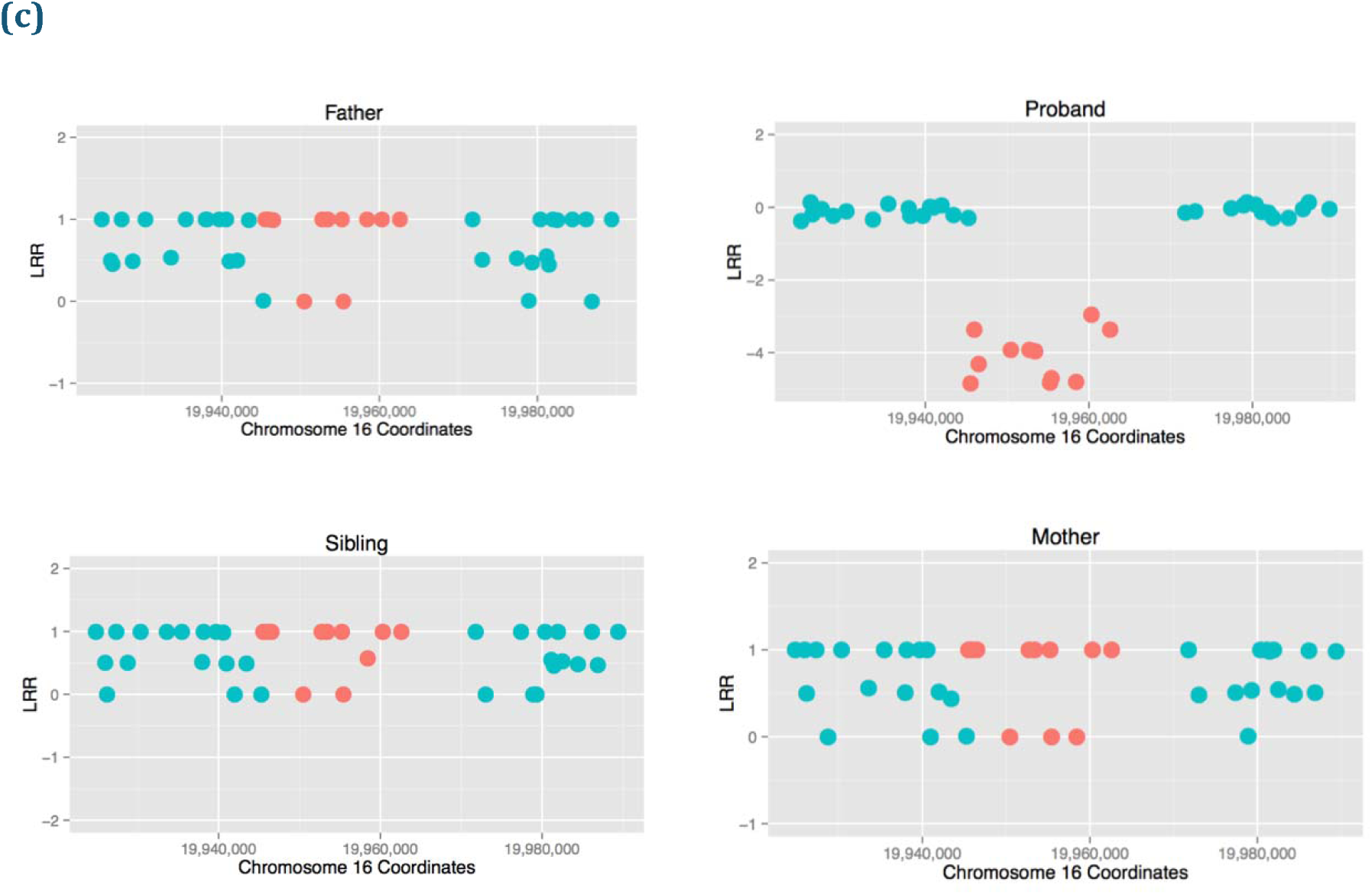
**(a)** Genome Browser Screen cut for the Read Depths in the K21 CNV 16p12.3 region of 22Kb. **(b)** B Allele Frequencies values for Illumina Omni 2.5 markers on K21 16p12.3 region including the markers belonging to the CNV region detected by ERDS in red. **(c)**. Log R Ratio values for Illumina Omni 2.5 markers on K21 16p12.3 region including the markers belonging to the CNV region detected by ERDS in red.

**Figure 7.**
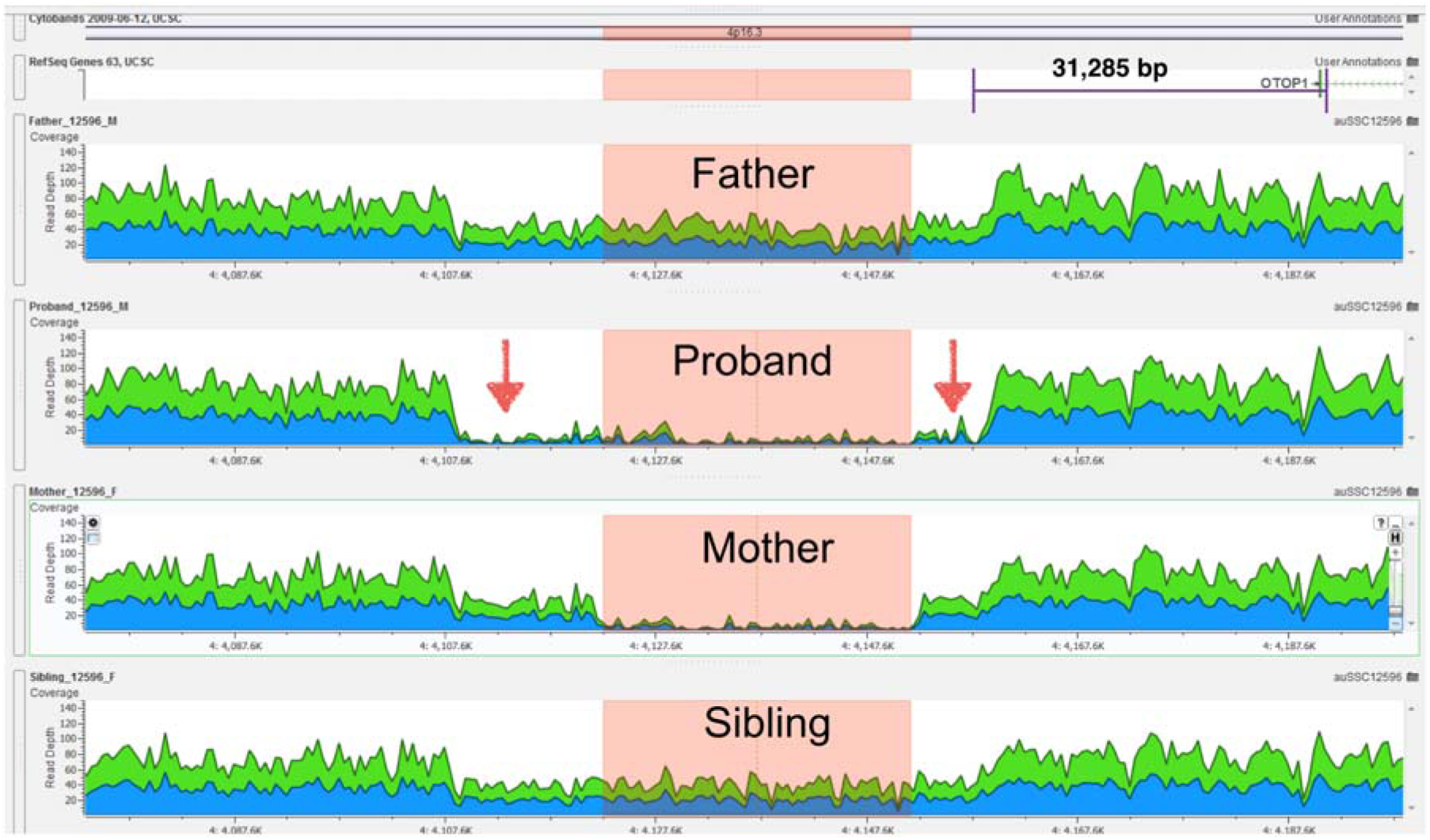
Genome Browser Screen cut for the Read Depths in the SSC_2 CNV 4p16.3 region of 50Kb. The highlighted region is where the 4 people bear either a homozygous or heterozygous deletion, only the Proband has an homozygous deletion of the complete deletion region, which could have been generated by inheriting the deleted copy from both parents.

**Table 6.** Number of variants obtained by each algorithm before and after filtering.

| Variants | GATK HC | Freebayes | Intersection | Unique to GATK % | Unique to Freebayes % | Intersection % |
| --- | --- | --- | --- | --- | --- | --- |
| Raw SNVs | 3,911,804 | 4,216,193 | 3,593,919 | 13.7 | 7 | 79.3 |
| Filtered SNVs | 3,403,728 | 3,714,842 | 3,255,217 | 11.9 | 3.8 | 84.3 |
| Raw Indels | 814,730 | 790,178 | 580,335 | 20.5 | 22.9 | 56.6 |
| Filtered Indels | 725,573 | 720,426 | 542,982 | 19.7 | 20.2 | 60.1 |

CGH microarray sequencing and analysis applied to the proband and his mother revealed the presence of a maternally inherited duplication spanning several genes (chrX:69074860-69512431). The duplication completely overlapped *OTUD6A*, *IGBP1*, *DGAT2L6*, *AWAT1*, *AWAT2*, *P2RY4*, *KIF4A*, *ARR3*, *GDPD2*, *RAB41*, *PDZD11* and partially overlapped *EDA*, *DLG3*, and *DLG3*. However, WGS-based CNV analyses revealed that the CNV was also present in the healthy sibling (Supp. Fig. 6a). PennCNV, which was applied to additional Illumina microarray data, did not accurately call the breakpoints for this CNV in any of the three individuals where it was initially detected (mother, proband, sibling), although its presence was clear from manual inspection of the microarray data (see Supp. Fig. 6b).

#### FMR1 test

Fragile X testing resulted in a normal number of CGG tri-nucleotide repeats for the K21 proband. Analysis of WGS data from all probands did not show any significant difference in CGG repeat content from the reference genome (Supp. Fig. 2-5). Traditional clinical Fragile X testing does not include sequencing *FMR1*, thus potentially missing other mutations that can contribute to the development of Fragile X syndrome [20-25]. Although the probands in this study did not present any of the common phenotypic features of Fragile X, a profile of all the CGG repeats present in every person was generated using WGS data and these profiles compared to the reference sequence (Supp. Fig 3-5). No point mutations reported in the literature as contributing to Fragile X were found in any of the probands [24].

#### Reproducibility of previous exome studies

As different approaches were taken to retrieve the variants for each proband, it was of interest to know if all of the SSC proband’s variants detected in the previous exome study, listed in Table 7 [13], were also detected by the methods used here. In cases where a variant was missed, this type of analysis will enable us to identify which step of the analysis pipeline might be responsible. Out of the three previously reported variants, which belong only to one family (SSC_2) only one was included in the final list of variants with this pipeline. Two of the three were lost by GATK after the initial filtering step, but they were still included in the downstream analysis because Freebayes and the Multinomial Analyzer still detected and retained them in their call sets. However, they were ultimately discarded after the CAAD score prioritization step, as they were not included in the top 1% most deleterious variants (<20 CAAD score). No variants were found in the SSC_1 family, and none of the variants reported in Table 7 were found in SSC_1 or K21.

**Table 7.** De Novo Indels.

| Algorithm | De Novo Indels |
| --- | --- |
| HC u FB | 52933 |
| Scalpel | 26 |
| Intersection | 4 |

## Discussion

### Concordance between algorithms

It is known that different algorithms are better at calling particular types of variants, each capable of detecting variants others cannot, and that they all usually agree on a subset of reliably called regions [26]. For this reason, the results of different algorithms were integrated, and instead of considering only the intersection of variants common to all algorithms, the union of all variant sets was obtained. This enabled the retention of variants that would have otherwise been excluded due to performing intersections with call sets, as only variants agreeing among all callers would have been retained. Indeed, one of the steps in which many variants are lost is during the initial filtering steps applied to each algorithm’s raw call set, at which point one can to decide how stringent the filter should be. Even recommended filtering parameters resulted in a detectable level of false negatives, i.e. true variants excluded from the final call set, despite these parameters being optimized for both sensitivity and specificity. Because the probands included in our study had already been part of previously reported targeted sequencing experiments, we were able to leverage available validation data to identify which informatics steps would have resulted in false negative calls. In our study, we found that for the GATK HC call set, not all of the previously validated calls [13] passed the first initial recommended filtering steps.

To measure concordance between the different variant calling algorithms used here, we considered variants in agreement if they match in terms of the genomic positions where each algorithm made a call. Due to large differences in INDEL calling and reporting, the same INDEL can sometimes be reported differently [27]. For this reason, the reference and alternative fields were not included in the analysis of concordance between the different INDEL callers. Another reason for comparing callers in this way is based on the large differences seen in multiallelic calls reported by GATK HC (∼30K) and Freebayes (∼70K). This non-standard way of reporting indels has made the comparison between algorithms particularly difficult, thus, the comparisons performed in this study are approximate. These issues underscore the importance of carefully integrating sets of variants from different variant callers, as simple intersections can dramatically reduce the number of true positives even if all callers detect them, as their representations may be slightly different between the different callers. New tools that standardize discordant variant reporting into a unified schema have been developed [28], and we expect that these tools will aid the in the more precise comparison and use of variants stemming from different callers.

### Microarray vs. WGS data for detecting CNVs

Microarray data provides researches with a cheap yet powerful way of detecting CNVs; however, depending on the particular technology used as well as the algorithms used to analyze the generated data, the results can vary widely. Sparse markers in some genomic regions makes it difficult to define accurate breakpoints of detected CNVs, something that is less difficult with CNVs detected from WGS data. This is due, in part, to the fact that WGS data is more uniform; having an average coverage of reads which is less variable across the genome. For this study we had both types of data for one of the quads, making it possible to call variants from both sources and compare the results. We found a large degree of variation in terms of the number of CNVs detected per person and also between the two detection methods used (that is, WGS based and DNA microarray based methods). The genome-wide sensitivity of CNV detection using WGS is higher, due to the fact that array based methods do not densely cover the entire human genome with markers. We found that having data from these orthogonal technologies was useful in including or excluding true or false positive calls, as each should show some evidence of a CNV, if one does exist. Thus, in regions where both technologies had enough data to detect CNVs, discordant calls could be easily resolved by comparing the data profiles between the two.

### Prioritization methods

Variant prioritization is another potentially delicate and important step in finding candidate disease contributing variants. One could detect all true variants from WGS analysis yet still discount biologically important variants if the pertinent annotations are not used correctly. When filtering based on annotations that are numerically scaled, filtering threshold values should generally be strict enough to result in a small number of variants in which functional studies are feasible, without letting any biologically important variant go unconsidered. Obtaining this variant set from a single annotation or score is currently not possible, as each individually lacks the power to filter to a small and manageable set, which could otherwise be obtained by using multiple annotations for threshold-based filtering. For these reasons, several tools and annotations were combined in order to make sure that the results were robust and not due to systematic errors from one prioritization framework. Although two different frameworks were used, they were only slightly different in their results, likely due to the fact that the VEP-GEMINI toolset has more annotations to determine if a variant is deleterious than does the in-house toolset. Unfortunately, using these two methods, we were unable to find a single candidate SNV or INDEL variant for the SSC_2 pedigree. One alternative would be to use other prioritizing methods, such as the Variant Annotation, Analysis, and Search Tool (VAAST), which employs an aggregative variant association test that combines both amino acid substitution (AAS) and allele frequencies and incorporates information about phylogenetic conservation [29]. As human variation not only includes small variations (SNVs and INDELs) and CNVs, but also structural variants and repeats, other software tools have to be used on these WGS data to explore other sources of variation that might contribute to disease.

### Candidate variants

After initial filtering, variant prioritization and segregation analyses, we found two *de novo* missense variants that were annotated as being highly deleterious (as defined by a CADD score of greater than 20) and rare on the population level (with population allele frequencies less than 0.01). The first variant was found in the proband of the SSC_2 pedigree and it is a stop gain variant in *MYBBP1A*, and the second is a missense in *LAMB3* found in the proband from the K21 pedigree.

### Stop gain in MYBBP1A

*MYBBP1A* codes for a nucleolar transcriptional regulator that was first identified by its ability to bind specifically to the Myb proto-oncogene protein [30]. The encoded protein is thought to play a role in many cellular processes including response to nucleolar stress, tumor suppression and the synthesis of ribosomal DNA, and many cancers have been previously associated with MYBBP1A including brain glioma [31]. According to UniProt [32], it may activate or repress transcription via interactions with sequence specific DNA-binding proteins and repression may be mediated at least in part by histone deacetylase activity. It has been shown that its down-regulation induces apoptosis and mitotic anomalies in mouse embryonic stem cells, embryonic fibroblasts and human HeLa cells [33]. The known information about MYBBP1A does not make any obvious connection to ASD, however it has not been possible to create a homozygous knock out mouse for MYBBP1A and this is thought to happen as it is essential for early mouse development prior to blastocyst formation [33]. In this study the mutation found in this gene is heterozygous and although healthy heterozygous knock out mice have been reported, it is not clear if those mice had similar any behavioral phenotypes related to autism; further studies are needed before any conclusions about the relevance of this variant in the etiology ASD can be made.

### LAMB3

*LAMB3* codes for a beta subunit laminin that belongs to a family of basement membrane proteins. Together with an alpha and a gamma subunit, LAMB3 forms laminin-5. It’s known that mutations in this gene can cause Autosomal-dominant Amelogenesis Imperfecta [34], epidermolysis bullosa junctional Herlitz type, and generalized atrophic benign epidermolysis bullosa, diseases that are characterized by blistering of the skin [35]. According to UniProt [31], its function is to bind to cells via a high affinity receptor, laminin is thought to mediate the attachment, migration and organization of cells into tissues during embryonic development by interacting with other extracellular matrix components. Again, the known diseases associated with this gene do not have an obvious link to autism, but its participation during embryonic development makes it an interesting candidate for further functional studies.

Although we found CNVs and SNVs that fit the filtering and annotation criteria described in the Methods section, there is no obvious connection between any of them and ASD, so they should be carefully considered only as possible disease contributing variants that are in need of further functional analysis. In addition, we did not have the statistical power of a larger study to be able to associate our variants as casual factors in the development of autism, and so our results are restricted to interpretation in the context of the three families studied here.

## Conclusions

Although a subset of ASD cases are now better understood, with their genetic contributions becoming more clear [2, 6, 8, 10, 13, 16, 17], the large degree of phenotypic heterogeneity in ASD leaves the vast majority of cases still poorly understood. Larger studies have focused on a subset of variant types, but here we have obtained a broader and more complete view of all the different types of genomic variation that could be contributing to the ASD phenotypes observed in this study. By combining different algorithms and variant prioritization methods, we were able to use the strengths of each and compensate for the different weaknesses by integrating their results in one computational framework.

There has been special interest in knowing to what extent *de novo* mutations are responsible for ASD cases [13]. In this study, four different variant detection algorithms and three different prioritization methods were used to detect *de novo* variants. This allowed us to improve detection sensitivity and to reduce the false negative rate. We also searched for variants segregating according to other disease transmission models, including autosomal recessive, X-linked, mitochondrial and compound heterozygote. As expected, the number of possible disease-contributing variants detected from each model varies widely (Table 4).

As sequencing technologies improve in accuracy and their operational costs decrease, large sequencing studies including thousands of people at higher sequencing depths are becoming more common. As such, it is useful and perhaps even necessary to design studies that search for and aim to detect all known variant types and to not just focus on a small subset. We suspect that such studies would, in general, obtain more biologically relevant results by doing so. However, study design must also consider the cost/benefit balance of sequencing whole genomes of a large number of people to high sequence depths, as was done here with the SSC quads (∼75X). The previously reported ideal coverage for accurately detecting SNVs is 40-45X where the detection saturates [36]. It has recently been shown that for accurate INDEL detection in personal genomes, whole genome sequencing coverage of 60X may be ideal, at least with 100 bp paired end reads from Illumina [37]. Given the known complexity and heterogeneity of ASD [2,6,9], it is clear that a large study capable of obtaining robust statistical signals is needed; yet a study of this magnitude with 60X coverage is still prohibitively expensive. Our study is useful in terms of contributing a small but rich dataset to larger studies, so that the etiology of ASD can be better understood. While this study was being completed, a study was published using the Complete Genomics (CG) platform to study 85 quartet families with autism [38], although there is a very high false negative rate associated with this sequencing technology, at least with the CG v2.0 pipeline [26].

## Methods

### Sample collection and sequencing

A pilot study of two SSC families and one Utah family was conducted. The Simons Simplex Collection (SSC) was assembled at 13 clinical centers, with the blood drawn from parents and children (affected and unaffected) sent to the Rutgers University Cell and DNA Repository (RUCDR) for DNA preparation. WGS was performed at CSHL on the two SSC families using the Illumina HiSeq 2000 platform at an average coverage of 75X, using paired-end 100 bp reads.

The Utah family had previously undergone fragile X screening and Chromosomal Microarray (CMA) genotyping for the proband and mother at the University of Utah. K21 blood samples were collected at the Utah Foundation for Biomedical Research, and genomic DNA was extracted and purified. Finally the DNA was quantified using Qubit dsDNA BR Assay Kit (Invitrogen) and 1 microgram was sent to the CSHL sequencing facility where WGS was performed on the Illumina HiSeq 2000 platform at an average coverage of 40X using paired-end 100 bp reads, and a parallel DNA samples was genotyped with an Illumina Omni2.5 array at the CHOP core facility.

### Fragile X analysis (FMR1 test)

The pedigree K21 proband was tested for Fragile X syndrome, a common inherited form of intellectual disability and autism spectrum disorder with characteristic phenotypic features, in which the majority of patients exhibit a massive CGG-repeat expansion mutation in *FMR1* that silences the locus [24]. In order to know if the expansion was present, the fragile X region was amplified by PCR using a single chimeric primer set in which one of the primers is fluorescently labeled. The reactions were then separated by capillary electrophoresis on the ABI310xl Genetic Analyzer and analyzed using the GeneMapper software. Fragile X syndrome can sometimes be misdiagnosed as autism in the absence of the CGG repeat expansion. There are two missense and other point mutations in the FMR1 gene that have been reported and described as causative of Fragile X Syndrome [20-25]. Because missense mutations cannot be detected using the CGG-repeat test and because WGS data was available for every proband, loci spanning *FMR1* were carefully analyzed to see if any of the probands had any possible disease contributing mutation (e.g., p.Ile304Asn, p.Gly266Glu, IVS10+14C→T and p.Ser27X). A CGG repeat analysis on the Fragile X region (chrX: 146,993,468-147,032,646, http://omim.org/entry/309550) was also performed for all the probands to confirm that the CGG repeat number was normal compared to the reference genome. This was done by calling variants and generating a gvcf file with the GATK Haplotype Caller software. The gvcf file contains all sites in the FMR1 gene, whether there is a variant present or not. Using the gvcf file, the gene sequences were inferred and each CGG tri-nucleotide was plotted as it appears within the FMR1 gene region, making evident any subtle difference in the amount or positions of the CGG repeats (Supp. Fig. 2). This simple method will only work if the CGG repeat size is covered by the read length of the sequencing technology used to sequence the samples, otherwise it would not align to the reference sequence. However if the reads are not long enough and few or no reads are aligned, we may still infer the presence of an expansion if there is an apparent deletion in the 5’ UTR of *FMR1*.

### Chromosomal microarray analysis (CMA)

The pedigree K21 proband was genotyped using the Affymetrix Cytogenetics Whole-Genome 2.7 Array, which has a total of 2,141,868 markers across the genome, including 1,742,975 unique non-polymorphic markers and 398,891 SNP markers. After finding a CNV with unknown pathogenicity on chromosome X, the mother was also genotyped using the same array to determine if the CNV was inherited.

### SNV and INDEL variant calling

Before proceeding to analyze the WGS data, the quality of raw sequencing reads was assessed using FastQC [http://www.bioinformatics.babraham.ac.uk/projects/fastqc/], which summarizes sequence quality metrics that can indicate whether there was a problem with the sequencing experiment. This quality control procedure is important, as the quality of the raw sequencing data needs to be assessed before performing further downstream analyses. As human genomic variation can range from single nucleotide changes to whole chromosome variations, different analyses need to be performed to retrieve most of the true variation present in each person. In this study, several software packages were used in an integrative manner to analyze all the data generated by the different high throughput technologies. Raw sequence read quality analysis was performed for all samples, followed by aligning them to the reference genome. All analysis prior to the use of variant caller software were applied to the data in a lane by lane fashion; this is done in order to take account for experimental variation introduced by optical duplicates known to occur in a lane specific manner [39].

### Whole genome sequence aligning and pre-calling processes

Whole Genome Sequence reads from all samples were aligned, lane by lane, to the GRCh37/hg19 human reference sequence using BWA-MEM 0.7.5a-r405 software [40] with default parameters, tagging shorter split hits as secondary for compatibility with Picard tools used downstream of the alignment. Samples from the SSC families were sequenced to a mean coverage of 75X, with 6 different lanes per sample used to achieve this depth. K21 family samples were sequenced to a mean coverage of 40X, obtained by using 3.5 lanes. The resulting alignments were converted to binary format, then sorted and indexed using SAMtools version 0.1.19-44428cd [41]. Duplicated reads were marked and read groups were assigned to each lane using Picard tools v1.84 [http://sourceforge.net/projects/picard/]. The GATK Indel realigner v3.0-0 was used to correct mapping artifacts that due to reads aligning to the edges of INDELs, may look like evidence for SNPs. The GATK Base Quality Score Recalibrator was also used to correct systematic errors of sequencing technologies [39, 42, 43]. Finally all lanes were merged by sample with Picard tools to generate a ready-to-use alignment.

### Variant detection for SNV and INDELS

After obtaining ready-to-use alignments, four different variant callers were used to analyze the WGS data for each individual in the three different families. SNVs and INDELs were called using the GATK Haplotype Caller, v2.8-1 and v3.0-0, with default parameters. GATK Haplotype Caller variants were filtered using the GATK variant quality score recalibration (VQSR) tool. The Freebayes variant caller v9.9.2-43-ga97dbf8 [44] was also used to call SNVs and INDELs on all individuals. Freebayes calls with a QUAL score of less than 30 or with less than 10 supporting reads were filtered out. To further support the detection of *de novo* calls, two other packages were used: Scalpel [45] in *de novo* mode for *de novo* INDEL detection and the Multinomial Analyzer (MA) [13], which implements a multinomial model that considers evidence stemming from all members of a quad to decide whether a call is a true de novo or not. Both Scalpel and the Multinomial Analyzer were used with default parameters and filtering thresholds for MA were set to denovo score>60 and Chi2Pval > 0.0001, as was used in the exome study in which both SSC families were previously analyzed [13]. Variants from the same sample coming from GATK and Freebayes were merged into a single vcf file for downstream analysis. All variants in the final set were visually inspected on the Golden Helix Genome Browser [Golden Helix GenomeBrowse® visualization tool (Version 2.0.4)]

### Variant classification and prioritization

The final set of high quality calls were divided into different models of inheritance, so that the way in which the mutations emerged and how they were possibly contributing to the condition could be interrogated. After obtaining model-specific subsets, the variants were annotated with a Combined Annotation Dependent Depletion (CADD) score, a metric that evaluates the deleteriousness of SNVs as well as INDELs variants in the human genome. CADD scores are generated by integrating multiple annotations, including PolyPhen and SIFT scores, into one metric by contrasting variants that survived natural selection with simulated mutations [46]. Those variants with a CADD score of greater than 20 were kept as potentially deleterious, and the number of reads supporting each variant was compared among all family members to decide whether a call was a false positive or not. All variants were further filtered using a MAF < 0.01 from the 1000 genomes project (Oct 2014). The final set of variants was annotated with in-house tools as well as the ANNOVAR software [47] using the UCSC [48] and RefSeq [49] gene tables, the SSC [50], Exome Variant Server [http://evs.gs.washington.edu/EVS/] and ClinVar databases [51] and the recently released ExAC database [http://exac.broadinstitute.org)].

### Models

There are several ways in which a disease-contributing genetic variant can be present in an individual. As we were not only interested in the variants, but also in their origin, they were divided into different models before prioritization.

### De Novo Model

*De novo* variants are those that emerge at some stage during the gametogenesis of one of the parents or embryogenesis of the child, so those mutations will be only present in the offspring and not the parents. Only those variants present uniquely in the proband and not in parents or unaffected sibling were kept for downstream annotation and analysis.

### X-Linked

Here only variants on chromosome X are considered. As all of the probands in this study are males, the only X chromosome copy they have comes from the mother, who by having two X chromosome copies could be masking the deleteriousness of a mutation, which is then expressed fully in the male offspring. All X chromosome variants present in the proband inherited from the mother but not present in the healthy sibling or father were kept for downstream annotation and analysis.

### Autosomal Recessive

In this model, a given variant is required to be present in both probands with one copy inherited by the mother and the second one from the father. The autosomal recessive variants found in the healthy sibling are also excluded.

### Compound Heterozygous

Sometimes a gene can bear two different heterozygous mutations; one in each chromosomal copy, affecting both copies of a gene but not with the same exact mutation, as is the case for the autosomal recessive model. For this set of variants, only those combinations of heterozygous mutations on the same gene and present in the proband were considered.

### Mitochondrial

In a similar fashion as chromosome X, it is well known that the mitochondrial DNA is passed from mother to offspring; however in this case, if a mutation is contributing to the disease the mother would also be affected so the only mitochondrial mutations considered are under the de novo model described above.

### VEP-GEMINI

The VEP (Variant Effect Predictor)-Genome Mining (GEMINI) [52] toolset is a framework for annotating and prioritizing genomic variants by different criteria. Built-in analysis tools were used to obtain variants characterized by different classifications: *de novo*, compound heterozygous, autosomal recessive and impact severity. The VEP-GEMINI toolset was used to get additional information about each variant, and to compare the results obtained with the model classifications and prioritizations performed with in-house tools. The criteria for keeping variants from each classification scheme were for variants to have a CADD score of greater than 20 or be annotated as having high impact severity for the proband.

### Variant calling for copy number variants

The Estimation by Read Depth with SNVs (ERDS) software [53] was used with default parameters to call CNVs from WGS data on each individual. It uses WGS data along with previously generated vcf files using the read depth and number of contiguous heterozygous and homozygous SNVs to call CNVs. Only calls with an ERDS score of > 300 were kept.

Additionally, CNVs were called with the microarray data from pedigree K21, which was genotyped with an Illumina Omni2.5 array and analyzed with the software package PennCNV [54]. For kilobase-resolution detection of CNVs, PennCNV uses an algorithm that implements a hidden Markov model, which integrates multiple signal patterns across the genome and uses the distance between neighboring SNPs and the allele frequency of SNPs. The two signal patterns that it uses are the Log R Ratio (LRR), which is a normalized measure of the total signal intensity for two alleles of the SNP and the B Allele Frequency (BAF), a normalized measure of the allelic intensity ratio of two alleles. The combination of both signal patterns is then used to infer copy number changes in the genome. Microarrays often show variation in hybridization intensity (genomic waves), which is related to the genomic position of the clones, and that correlates to GC content among the genomic features considered. For adjustment of such genomic waves in signal intensities, the cal_gc_snp.pl PennCNV program was used to generate a GC model that considered the GC content surrounding each Illumina Omni2.5 marker within 500kb on each side (1Mb total). The joint-calling algorithm designed for parents-offspring trios was used, as it is the most accurate of the algorithms in the package for family based studies. The Hidden Markov Model used is contained in the hhall.hmm file provided by the latest PennCNV package, and the custom Population Frequency of B allele (PFB) file for all the SNPs in the Illumina Omni2.5 array was generated from 600 controls (which consists of 600 unaffected parents from the Simons Simplex Collection (provided by Dr. Stephan Sanders from Yale University). The GC model described above was also used during CNV calling.

Chromosome X CNVs were called separately using the -- test mode with the --chrx option. Using BEDtools [55] and in-house tools, consensus CNV calls were obtained for parents from the two separate trio calling processes that had to be done for each child in the quad. CNVs were quality filtered by considering the length of the CNV event (for both algorithms: ERDS and PennCNV) and for microarray data, the number of SNPs embedded on the CNV region and the number of expected SNPs for that given region (Supp. Fig. 5), histocompatibility regions, centromeric and telomeric regions were also filtered out as it is common to find non-pathogenic variants there (both algorithms).

For Pedigree K21, ERDS and PennCNV calls were compared and the union of each pipeline’s set of variants was annotated with in-house tools and the ANNOVAR software [47] using dbVar [56], DGV [57], ClinVar [51], DECIPHER [58], ENCODE [35] and the SFARIgene databases [50] and those variants which > 90% of their total length overlapped reciprocally with variants found in controls were ruled out. ERDS filtered output for pedigrees SSC_1 and SSC_2 were annotated with the same software and criteria.

### Sanger Sequencing

PCR primers for the Chr17:4458481(hg19) variant in *MYBBP1A* were designed to produce a 911 bp amplicon, using Primer 3 (http://primer3.sourceforge.net). Primers were obtained from Sigma-Aldrich®, and tested for PCR efficiency with an in-house DNA sample using a Phusion Flash High-Fidelity PCR Master Mix (Life Technologies, USA). The optimized PCR reaction was then carried out on patient DNA. PCR products were visually inspected for amplification efficiency using agarose gel electrophoresis, and were purified using the QIAquick PCR Purification Kit (QIAGEN Inc., USA). Purified products were then diluted to 5∼10 ng/μl in water for use with the ABI 3700 sequencer. The resulting *.ab1 sequence files were loaded into the CodonCode Aligner V5.1.2 for analysis. All sequence traces were manually reviewed to ensure the reliability of the genotype calls.

## List of abbreviations used

(SNP): Single-nucleotide polymorphism
(CNV): copy number variation
(INDELs): insertions and deletions
(SV): structual variant
(WGS): whole genome sequencing
(WES): whole exome sequencing
(NGS): next-generation sequencing
(base pair): bp
(kilo base pairs): Kb
(megabase pair): Mb
(polymerase chain reaction): PCR

## Additional Information

### Data Deposition and Access

All of the sequence reads can be downloaded under project accession number

[PRJNA282537] from the Sequence Read Archive (http://www.ncbi.nlm.nih.gov/sra).

SRA Bioproject: PRJNA282537

Biosamples: SAMN03571202, SAMN03571214, SAMN03571217, SAMN03571219

### Online Resources

1000G database: http://www.1000genomes.org/

Exome Aggregation Consortium (ExAC): http://exac.broadinstitute.org/

### Ethics compliance

Research was carried out in compliance with the Helsinki Declaration. Two of the families analyzed in this study belong to the SSC (referred as SSC_1 & SSC_2), and both families were clinically evaluated and extensively phenotyped as well as whole exome sequenced for a previous study [13].

The third family (referred to as K_21) was recruited to this study at the Utah Foundation for Biomedical Research (UFBR) where extensive clinical evaluation was performed. Written consent was obtained for phenotyping and whole genome sequencing through Protocol #100 at the Utah Foundation for Biomedical Research, approved by the Independent Investigational Review Board, Inc.

## Acknowledgments

Dr. Stephan Sanders provided the PFB file necessary for CNV calling. Dr. Kai Wang assisted in the CNV analysis. The authors would like to thank the Exome Aggregation Consortium and the groups that provided exome variant data for comparison. A full list of contributing groups can be found at http://exac.broadinstitute.org/about.

### Author Contributions

Jiménez-Barrón L.T. developed the study design, performed all informatics analyses and wrote the manuscript. O’Rawe J. provided in-house tools, contributed to the study design and revised the manuscript. Yiyang Wu performed the sanger sequencing validation experiment. H. Fang participated in the study design. Iossifov I. provided the SSC data. G.J.L. participated in the study design, data interpretation, and manuscript writing.

### Funding

The laboratory of G.J.L. is supported by funds from the Stanley Institute for Cognitive Genomics at Cold Spring Harbor Laboratory (CSHL). The CSHL genome center is supported in part by a Cancer Center Support Grant (CA045508) from the NCI.

### Competing Interests

G.J.L serves on advisory boards for GenePeeks, Inc. and Omicia, Inc.

---

**Table 8.** Final set of SNV.

| Model | Ref -> Alt/<br>Effect | Location hg19 | Affected<br>Gene | Algorithms<br>that called<br>the variant | Pedigree<br>ID | ExAC<br>Allele<br>Frequency | CADD<br>score |
| --- | --- | --- | --- | --- | --- | --- | --- |
| De Novo | Sub(C->T)<br>missense | Chr1:209823359 | LAMB3 | Freebayes<br>Multinomial<br>Analyzer<br>GATK | SSC_2<br>(12605) | 0 | 22.7 |
| De Novo | Sub(G->A)<br>nonsense | Chr17:4458481 | MYBBP1A | Freebayes<br>Multinomial<br>Analyzer<br>GATK | K21 | 1/74014=0.<br>00001351 | 40 |

**Table 9.** Previous SSC Exome Study Comparison.

| Family<br>ID | Location | Ref -> Alt/<br>Effect | Genes | Population<br>type | GATK_HC/<br>Filtered | Freebayes/<br>Filtered | MA/<br>Filtered | CAAD<br>Score |
| --- | --- | --- | --- | --- | --- | --- | --- | --- |
| 12605 | 10:103908608 | sub (C->T)/<br>missense | PPRC1 | denovo | yes/NO | yes/yes | yes/yes | 19.5 |
| 12605 | 1:209823359 | sub (A->C)/<br>missense | LAMB3 | denovo | yes/yes | yes/yes | yes/yes | 22.7 |
| 12605 | 3:185993461 | sub (G->T)/<br>intron | DGKG | denovo | yes/NO | yes/yes | yes/yes | 7.5 |

**Supplemental Figure 1.**
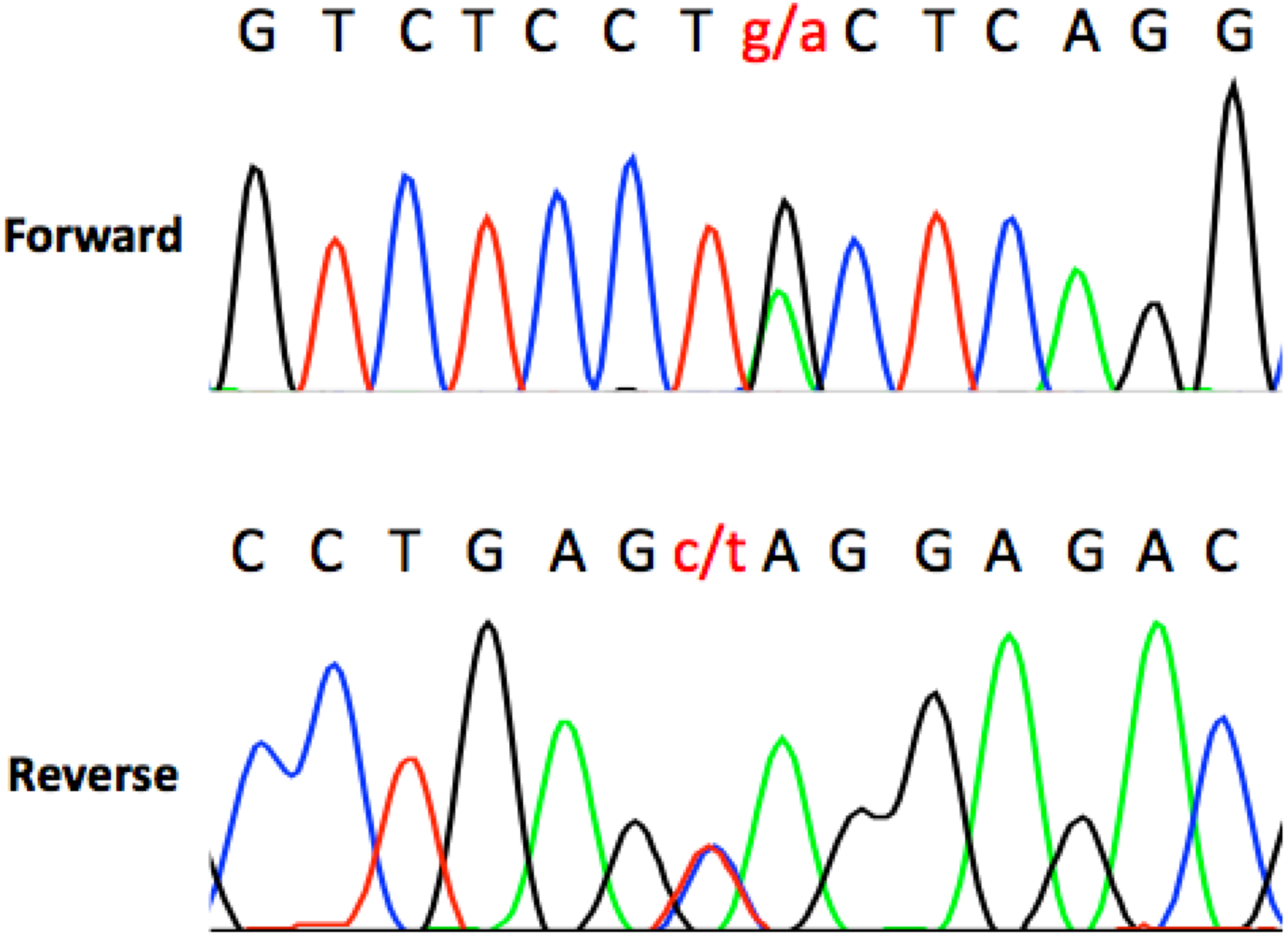
MYBBP1A stop gain validation by Sanger sequencing. Sanger sequencing validation shows two overlapping peaks: one for C and one for T on the reverse strand.

**Supplemental Figure 2.**
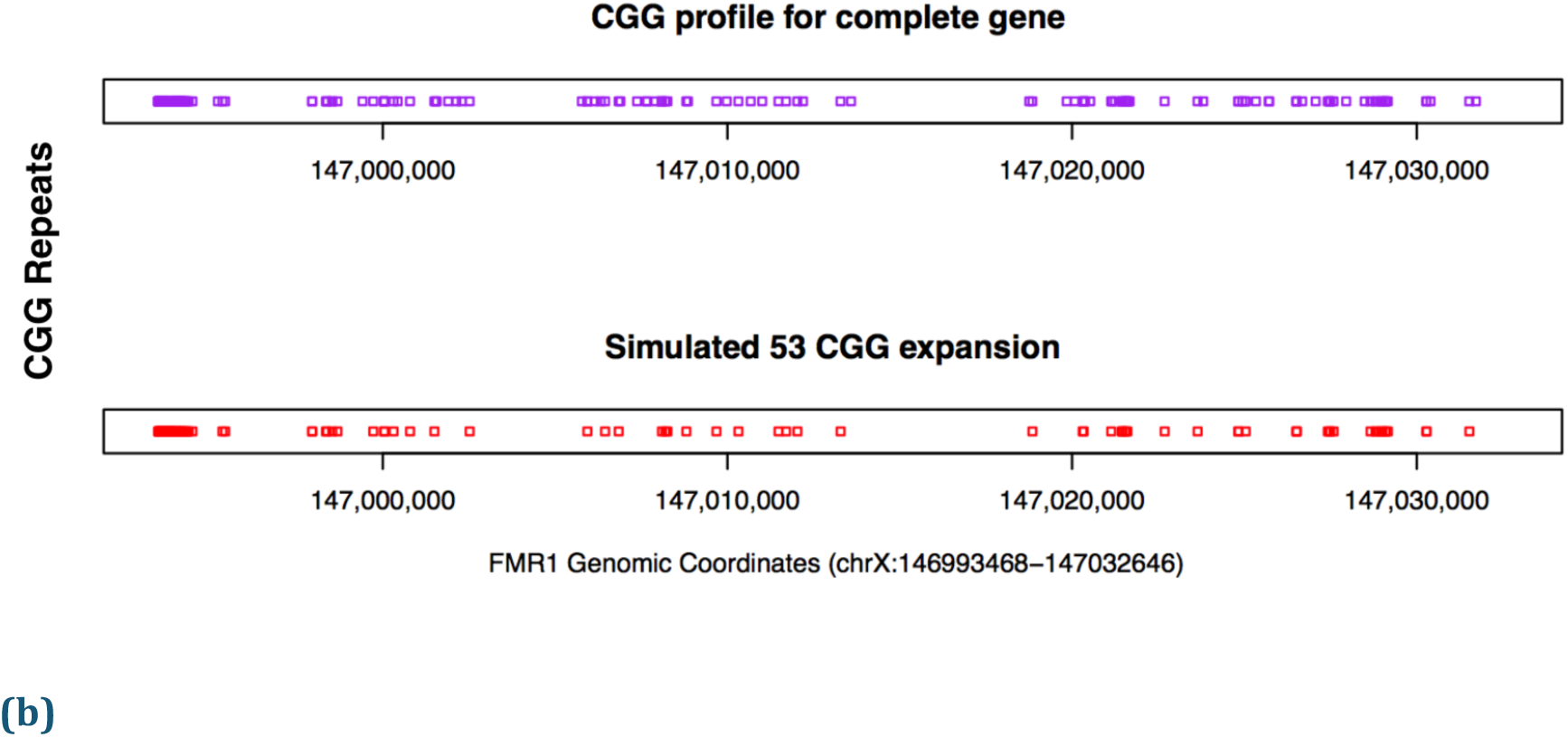
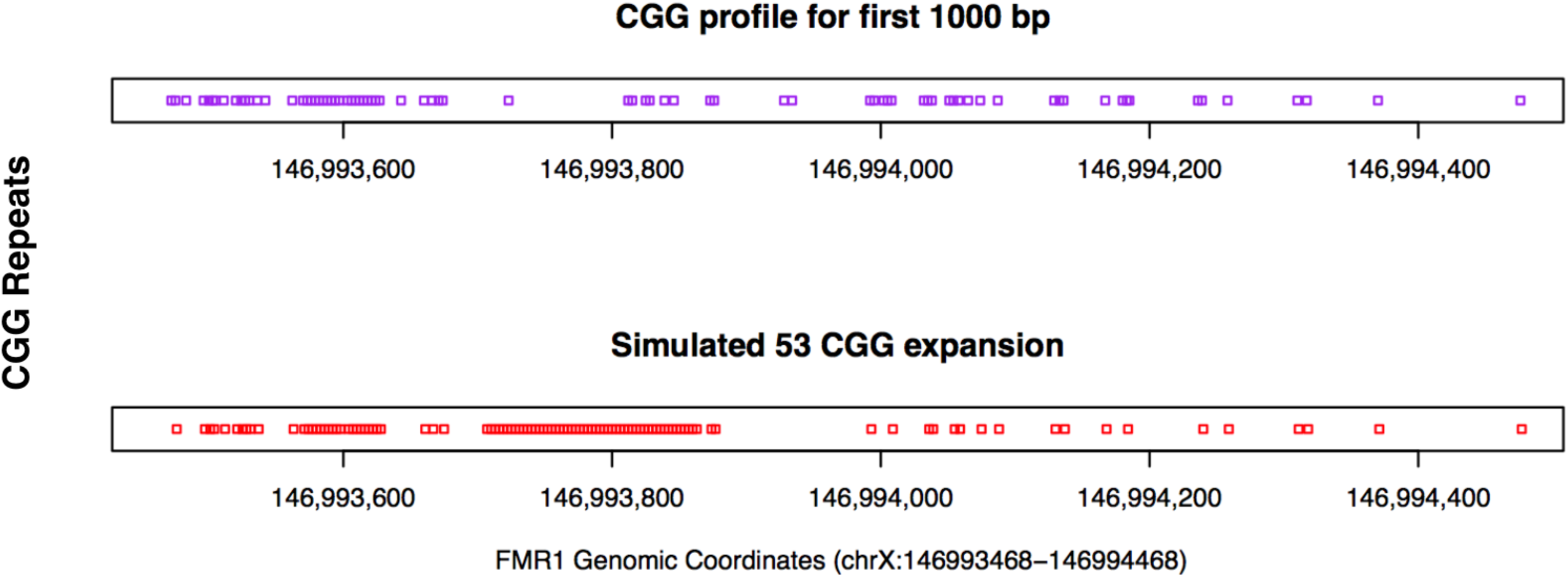
GGG repeats profile on the reference FMR1 complete gene and a simulated expansion. **(a)** The x axis represent the coordinates of the reference FMR1 gene, which includes the 5’UTR region where the CGG expansion occurs. **(b)** Here only the first 1000 closest nucleotides to the 5’UTR are plotted so a simulated expansion of randomly introduced CGG repeats is clearly appreciated.

**Supplemental Figure 3.**
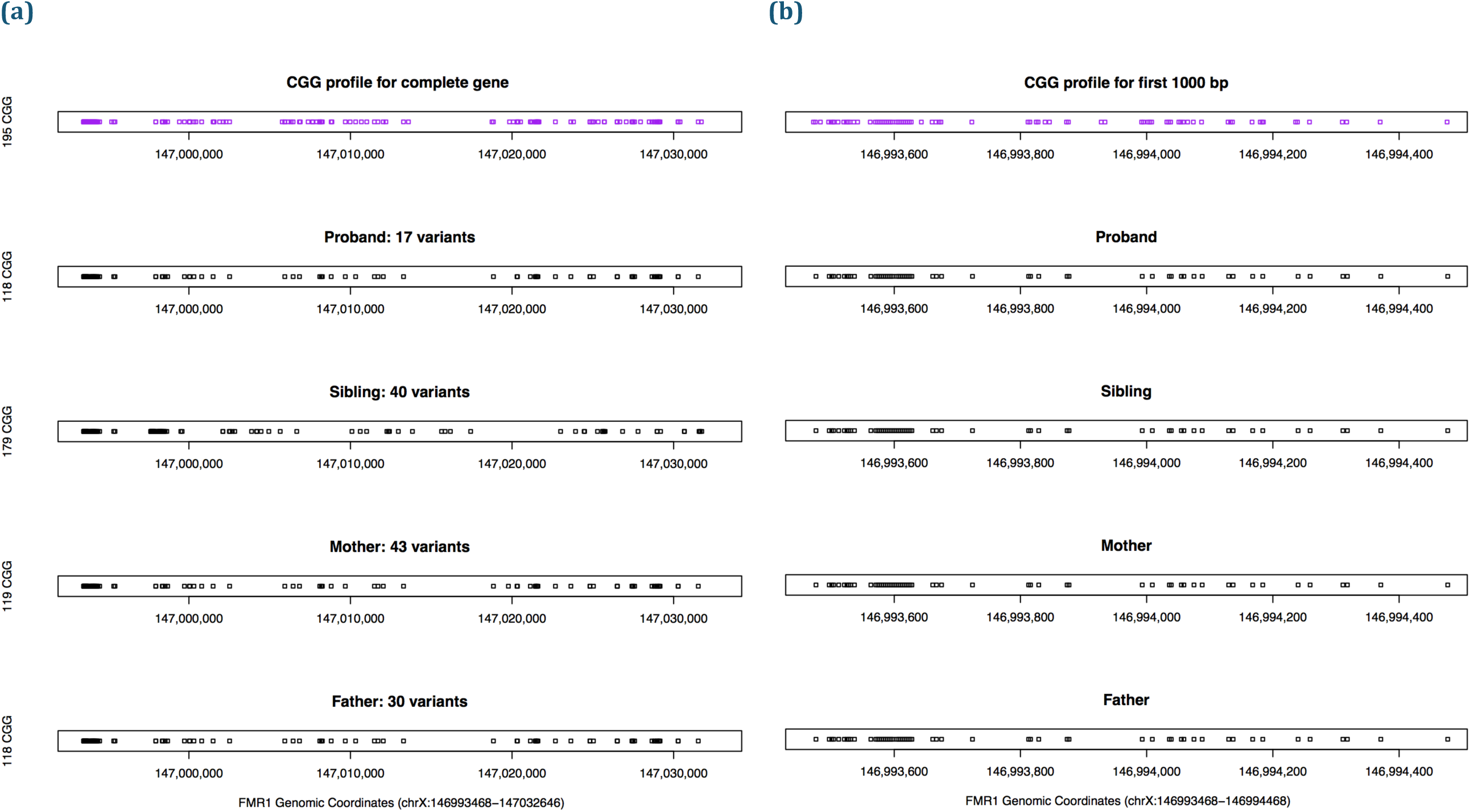
GGG repeats profile. Family K21 Complete FMR1 region profile. **(a)**The complete FMR1 gene CGG profile for this family looks normal, the number of CCG repeats for the proband is even less than the reference **(b)** First 1000 closest nucleotides to the 5’UTR.

**Supplemental Figure 4.**
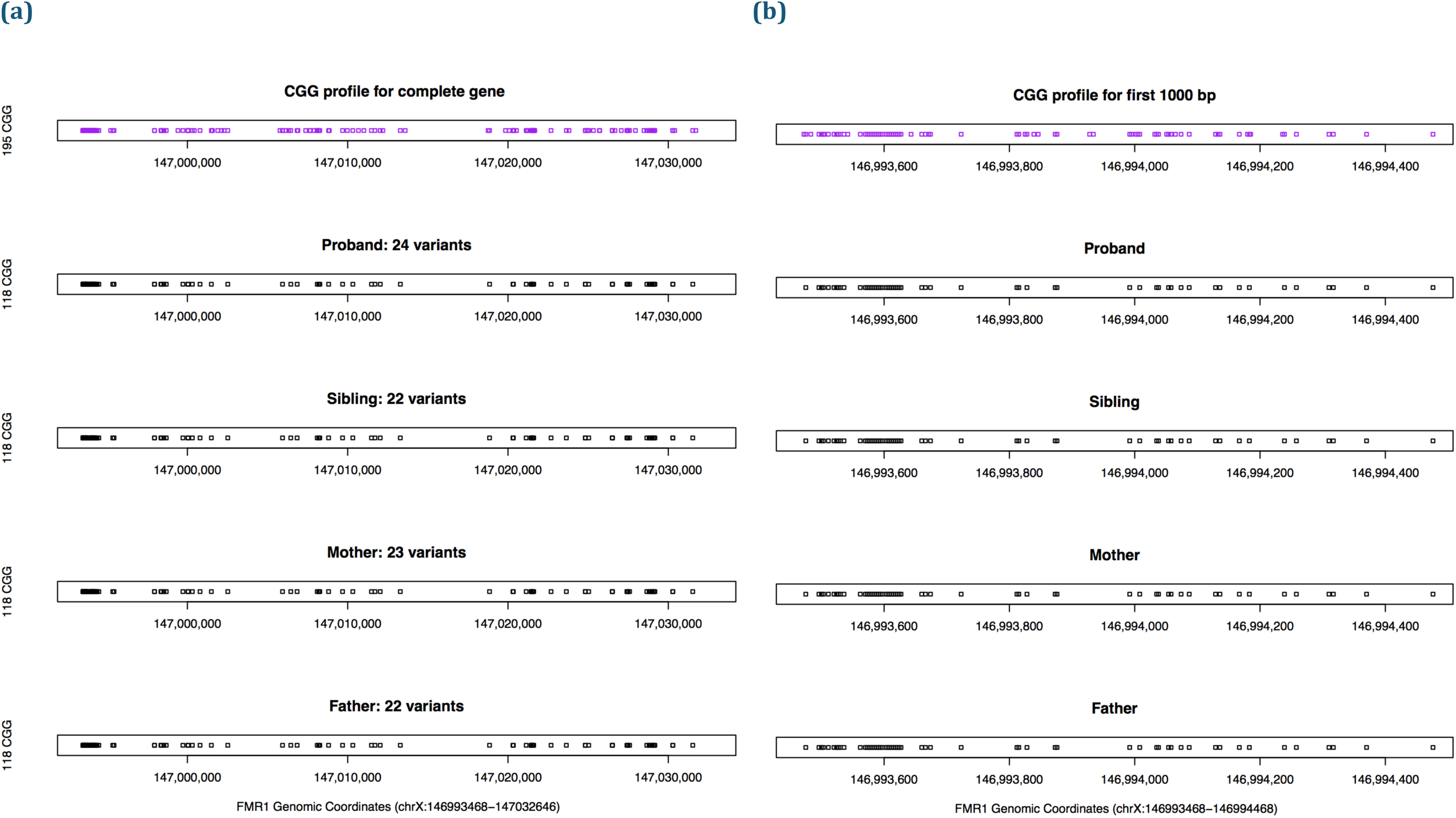
GGG repeats profile for family SSC_1. **(a)** The complete FMR1 gene CGG profile for this family looks normal. **(b)** The first 1000 closest nucleotides to the 5’UTR look normal.

**Supplemental Figure 5.**
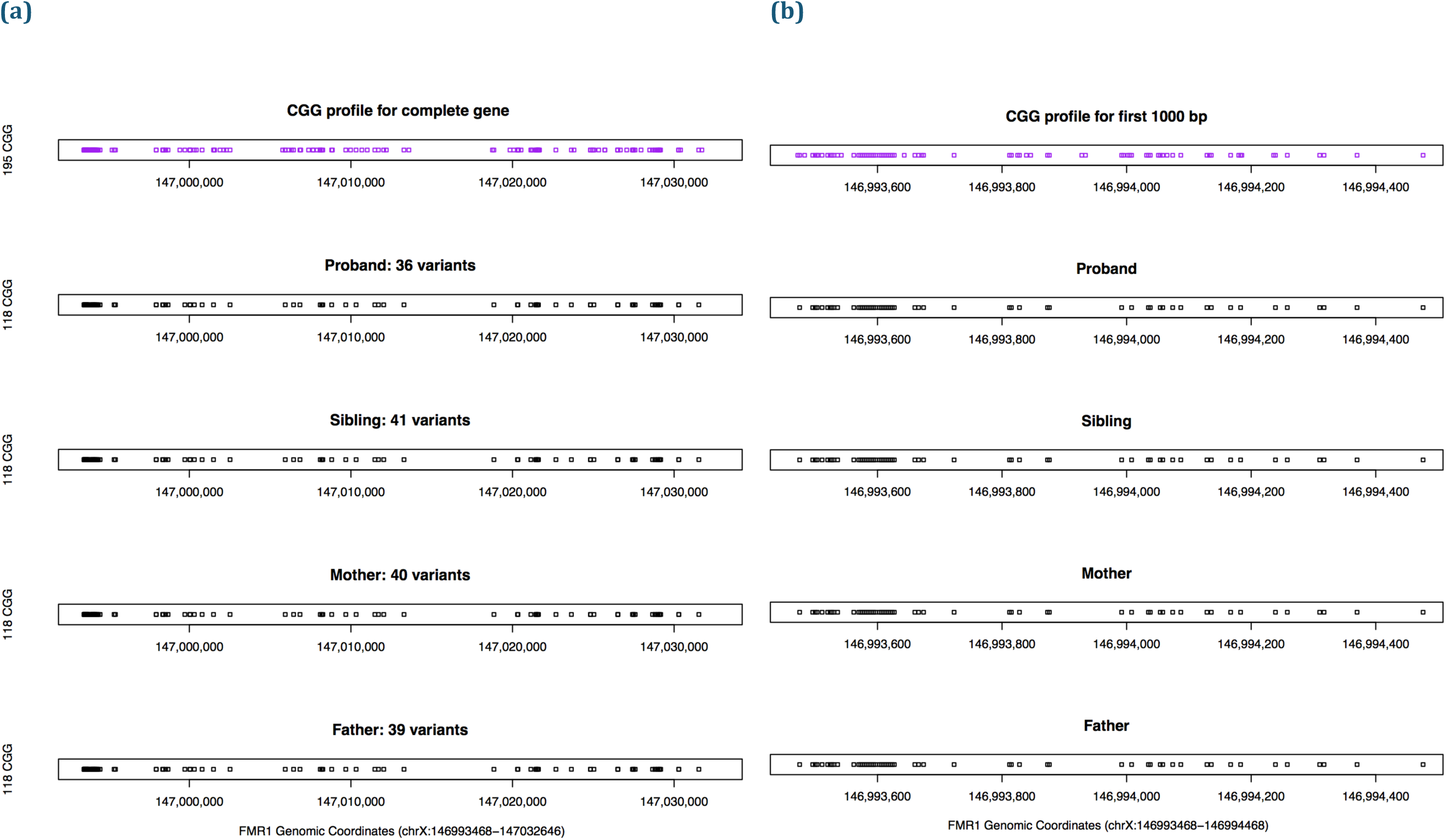
GGG repeats profile for family SSC_2. **(a)**The complete FMR1 gene CGG profile for this family also looks normal. **(b)** The first 1000 closest nucleotides to the 5’UTR look normal.

**Supplemental Figure 6.**
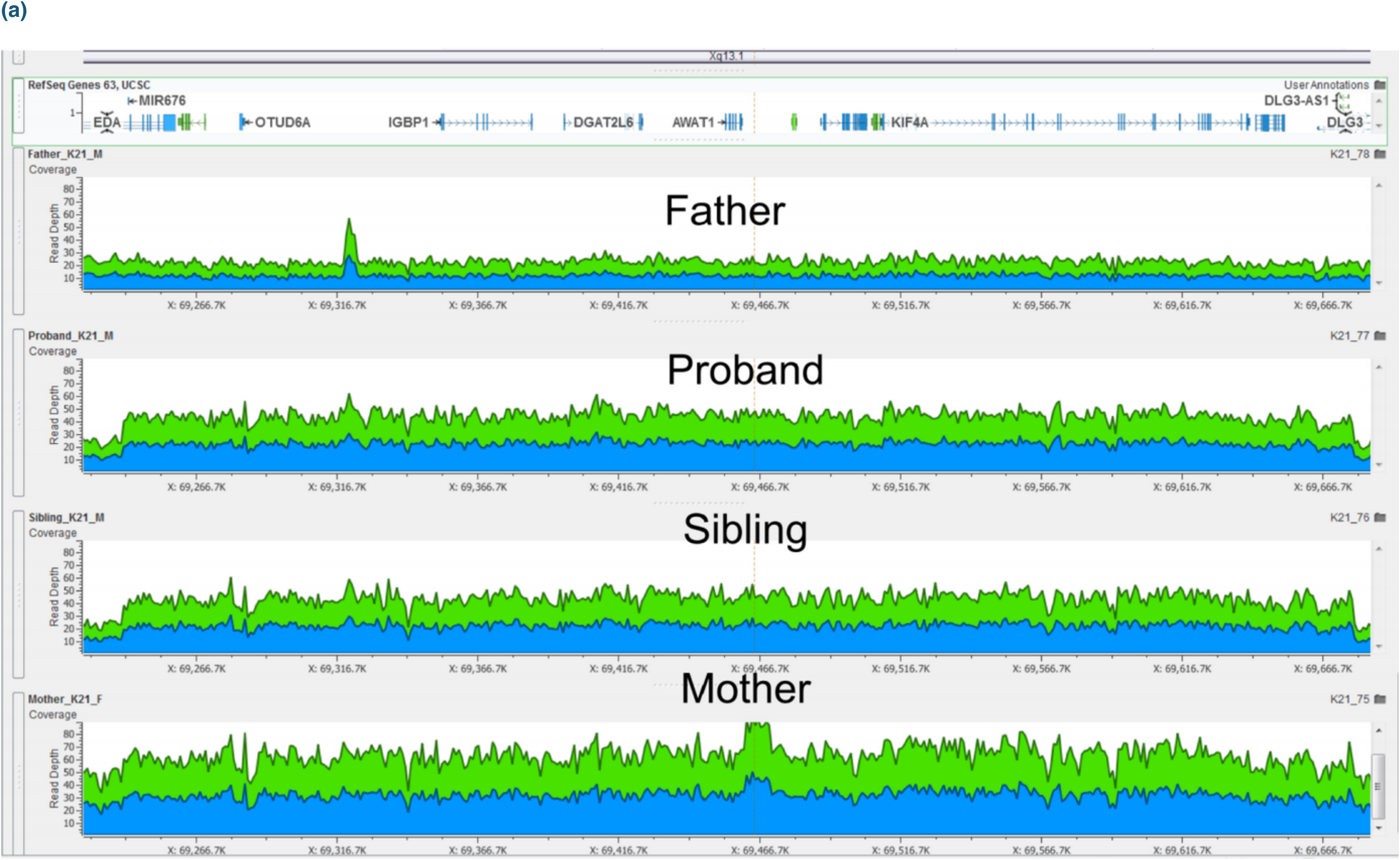
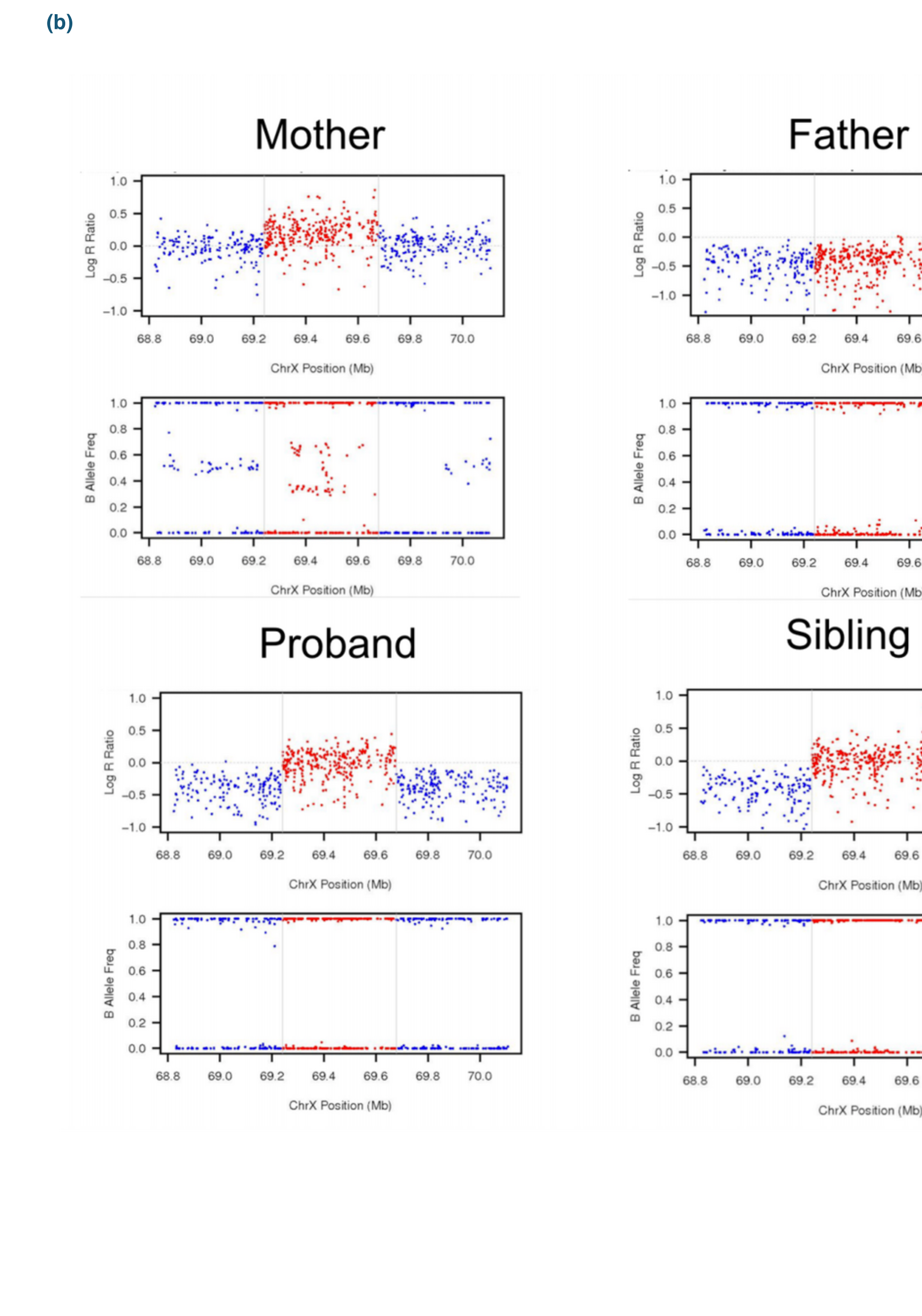
(a). Copy number variation on K21 detected by CMA, Illumina Omni2.5 array and WGS technologies. Here is the duplication region that was previously reported on the medical records which reported the following genes fully contained in the CNV: OTUD6A/IGBP1/DGAT2L6/AWAT1/AWAT2/P2RY4/KIF4A/ARR3/GDPD2/RAB41/PDZD11 and the following genes partially contained in the CNV: EDA/DLG3/DLG3 which actually corresponds to the genes contained in the CNV region reported by ERDS. As the healthy male sibling also inherited**. (b)** Even though PennCNV did not detect the CNV, by plotting the LRR and BAF values for all the family, the CNV can be confirmed to be present in the mother, proband and healthy sibling.

**Supp. Figure 7.**
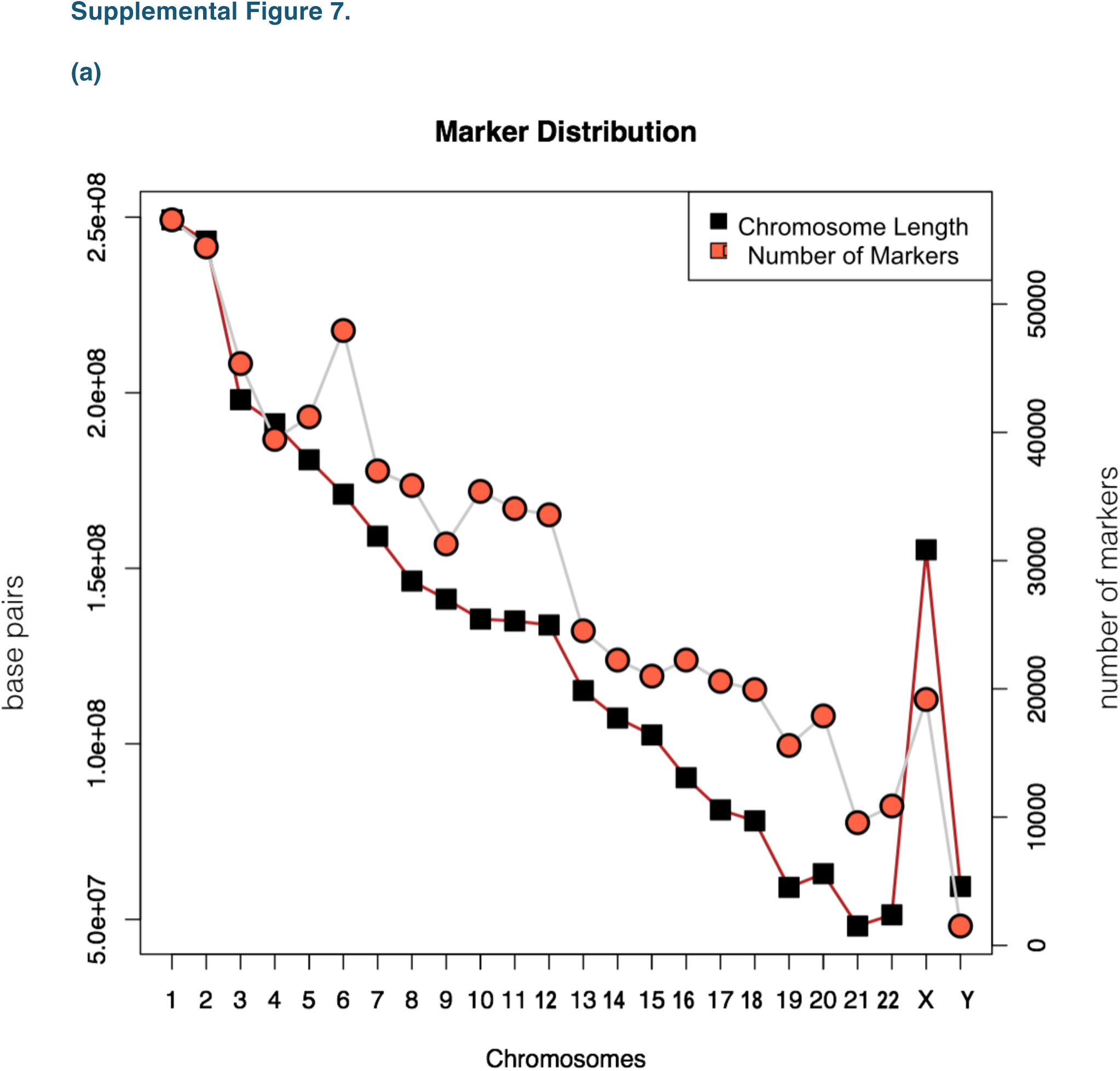
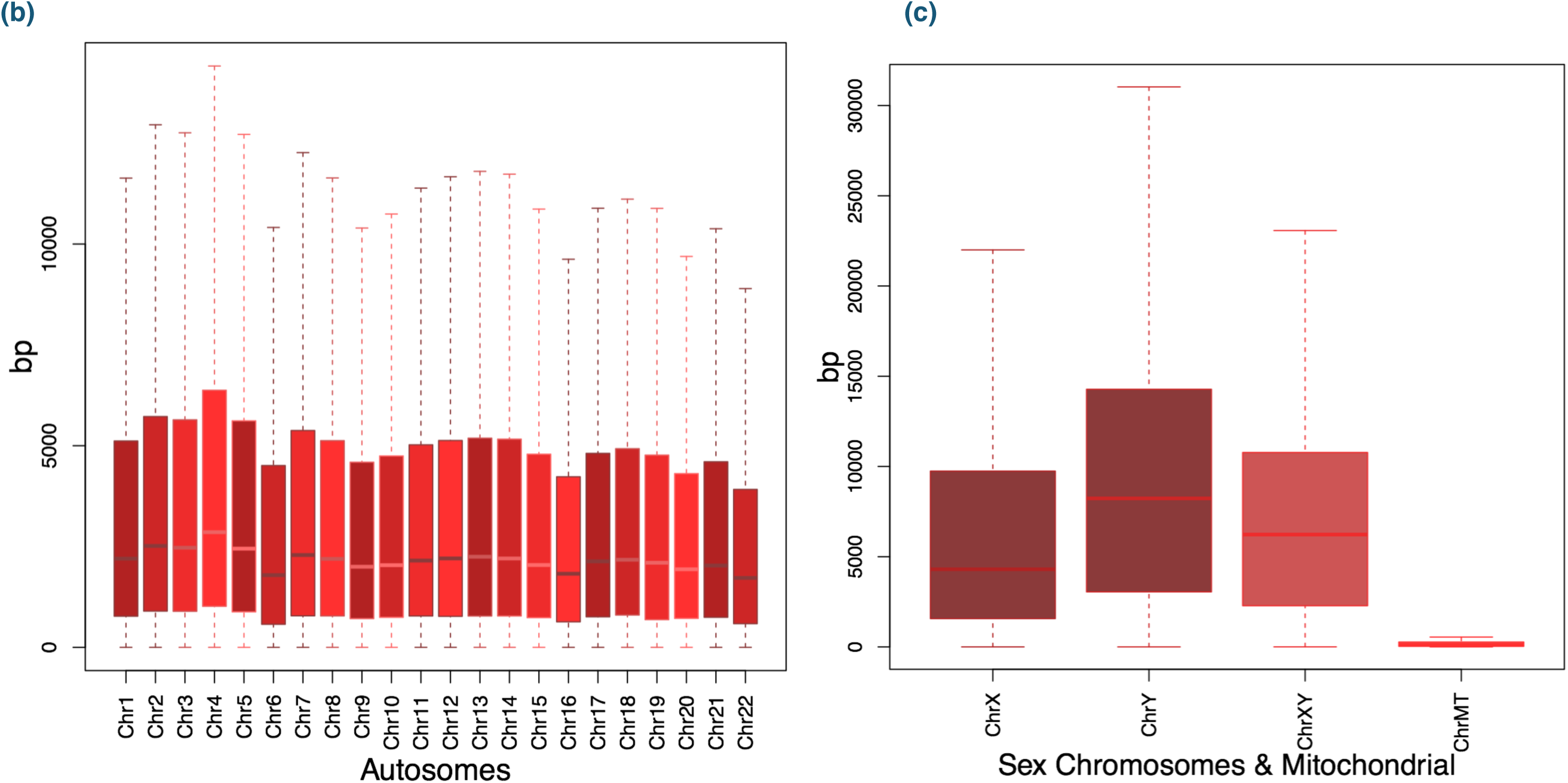
Markers Number by chromosome and the inter-marker spacing. The recommended methods for filtering Copy Number Variants called from microarray data are too arbitrary in the sense they are not aware of some features array-specific that could make this general filtering criteria suitable for all microarray calls. The number of markers and the space between them was evaluated for each chromosome and considered into the CNVs filtering criteria. **(a)** Here, the size of the chromosomes in base pairs (black) and the number of markers (red) are plotted, a line is drown across the dots so it’s easier to see that the greater the chromosome size, the greater the number of markers. However, the relation is not perfect and the number of markers for similar sized chromosomes, can vary largely. Because of this, the expected number of SNPs involved in a CNV call from one chromosome has to differ from those called in another chromosome. **(b)** However, not only the number of markers plays an important role but it was also important to make sure that the marker distribution across each chromosome was homogeneous without clusters of markers and large empty regions. If the size in base pairs of each chromosome was divided between the number of markers, the average inter-marker spacing should be 5Kb, to know if this was true, all the values for the spaces between markers (without outliers) are plotted here as quartiles, showing that 3/4 of the spacing values are around 5 or 6Kb and the other quarter having inter-marker spaces up to 10Kb, which explains why some regions are more difficult to call. **(c)** The sex chromosomes are plotted separately as their upper quartile has greater values than the autosomes. In the other hand, the Mitochondrial chromosome inter-marker spacing is smaller, this make sense as it’s size is only of ∼16Kb and the Illumina Omni2.5 microarray has 288 markers for it.

**Supp. Figure 8.**
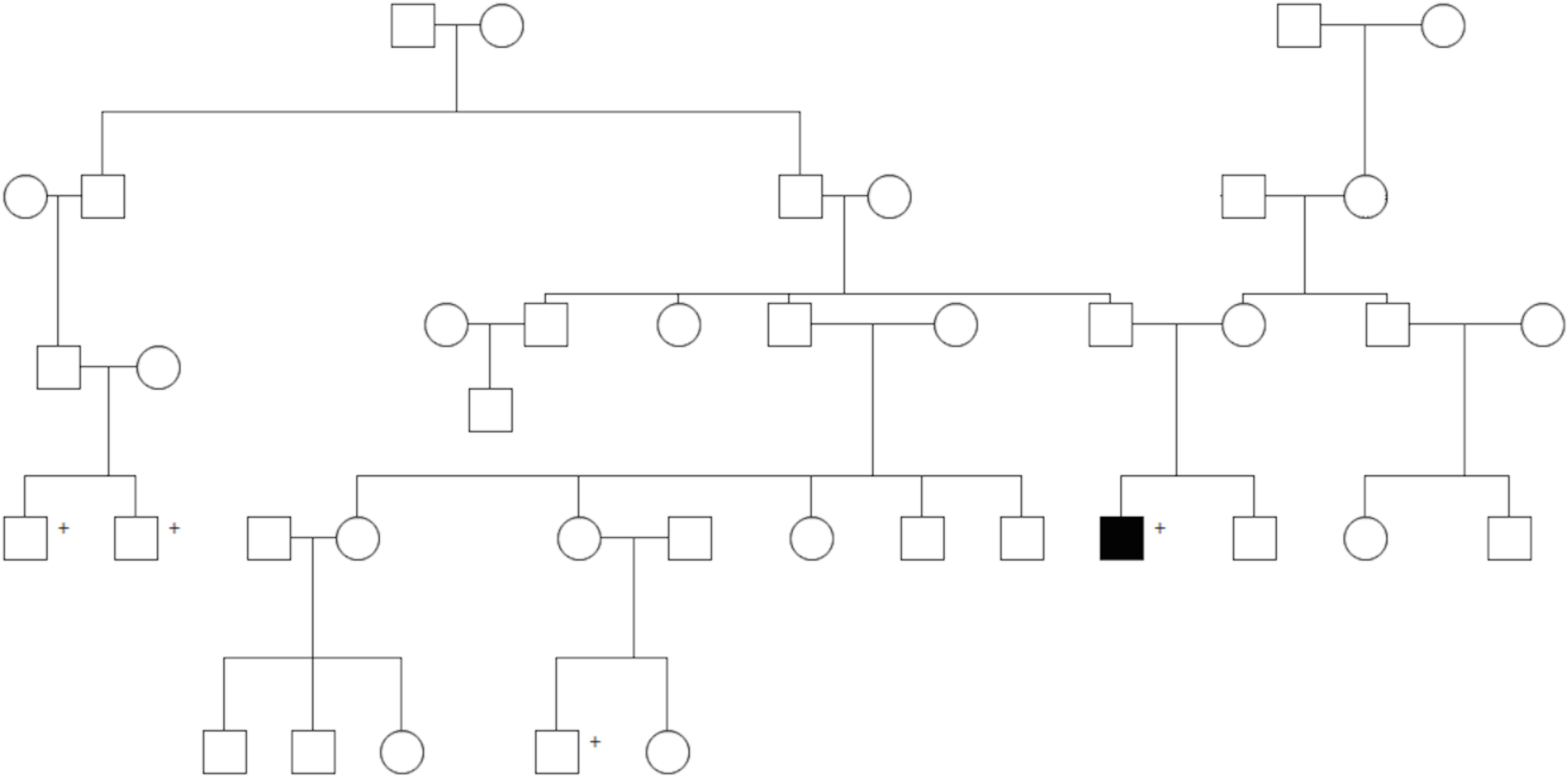
Extended Pedigree K21. Individuals with a + sign are affected by ???. The black square represents the K21 proband analyzed in this study.

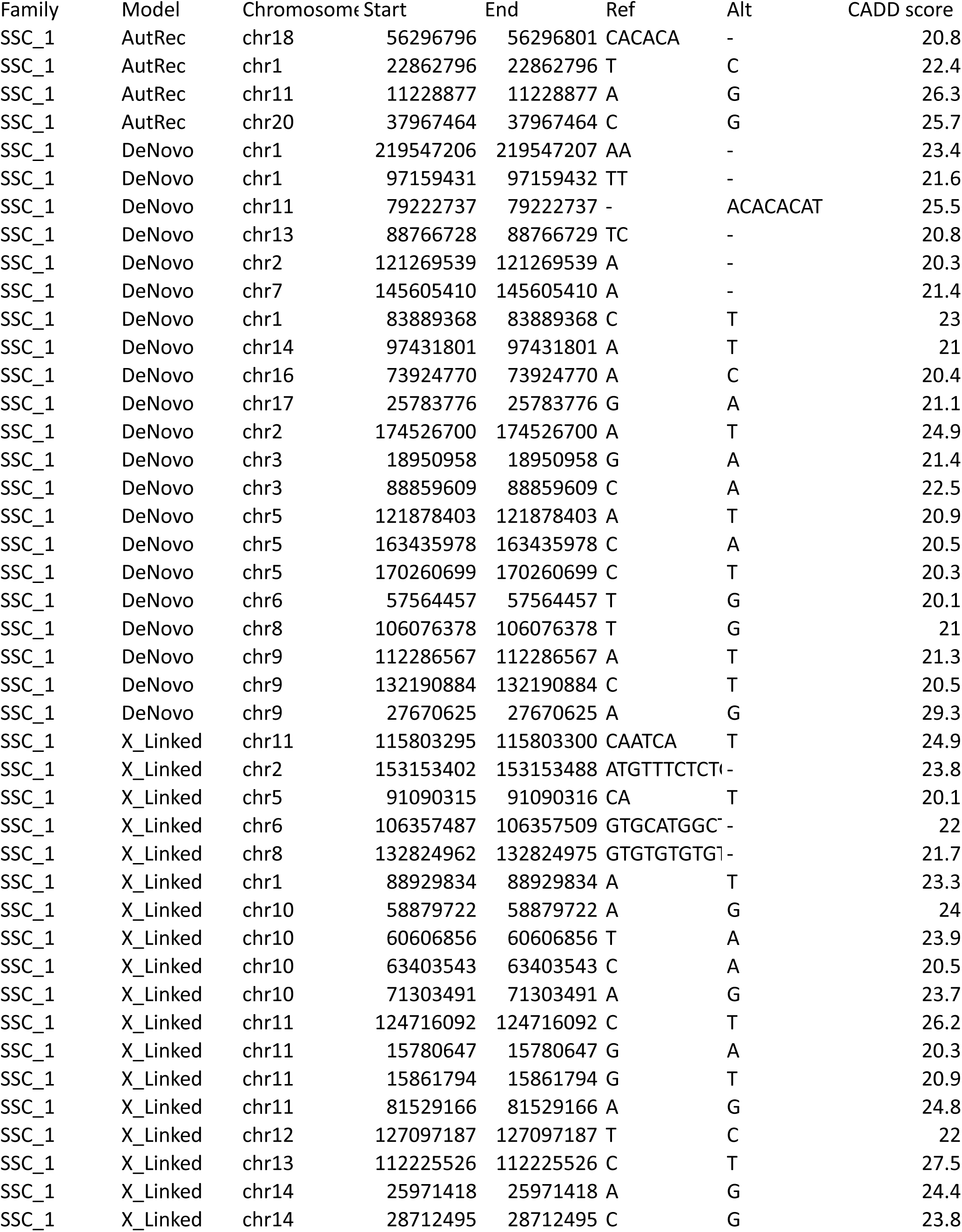
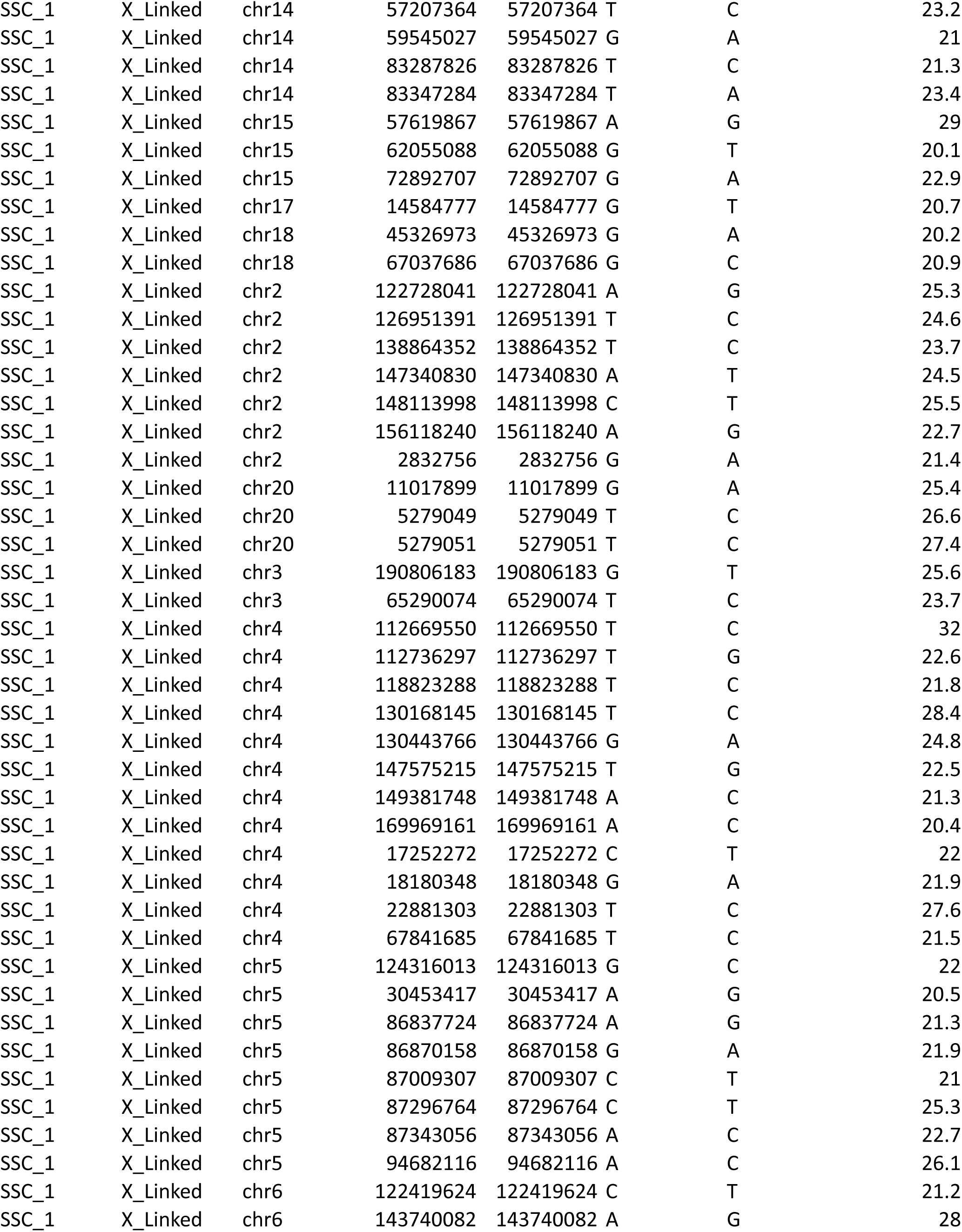
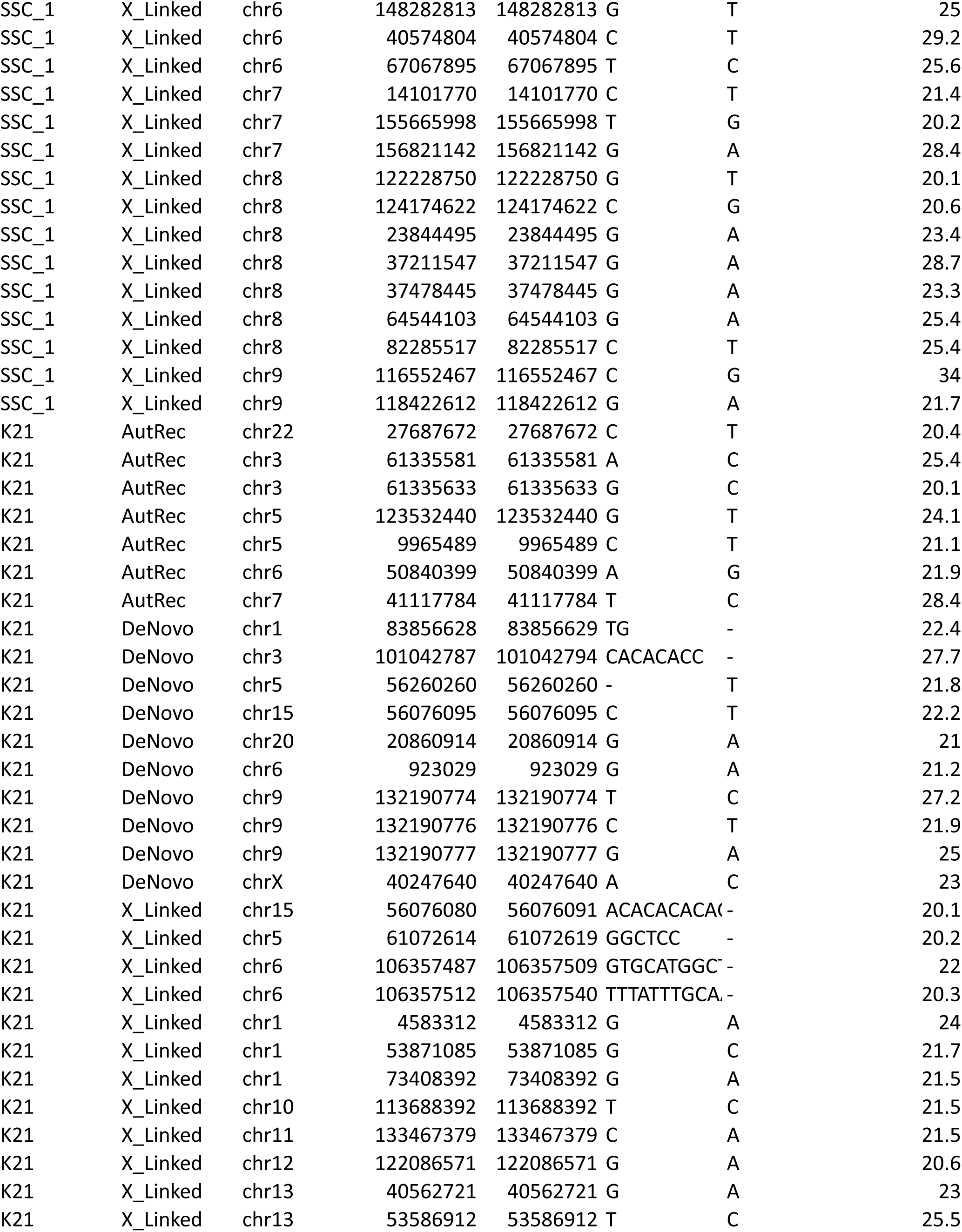
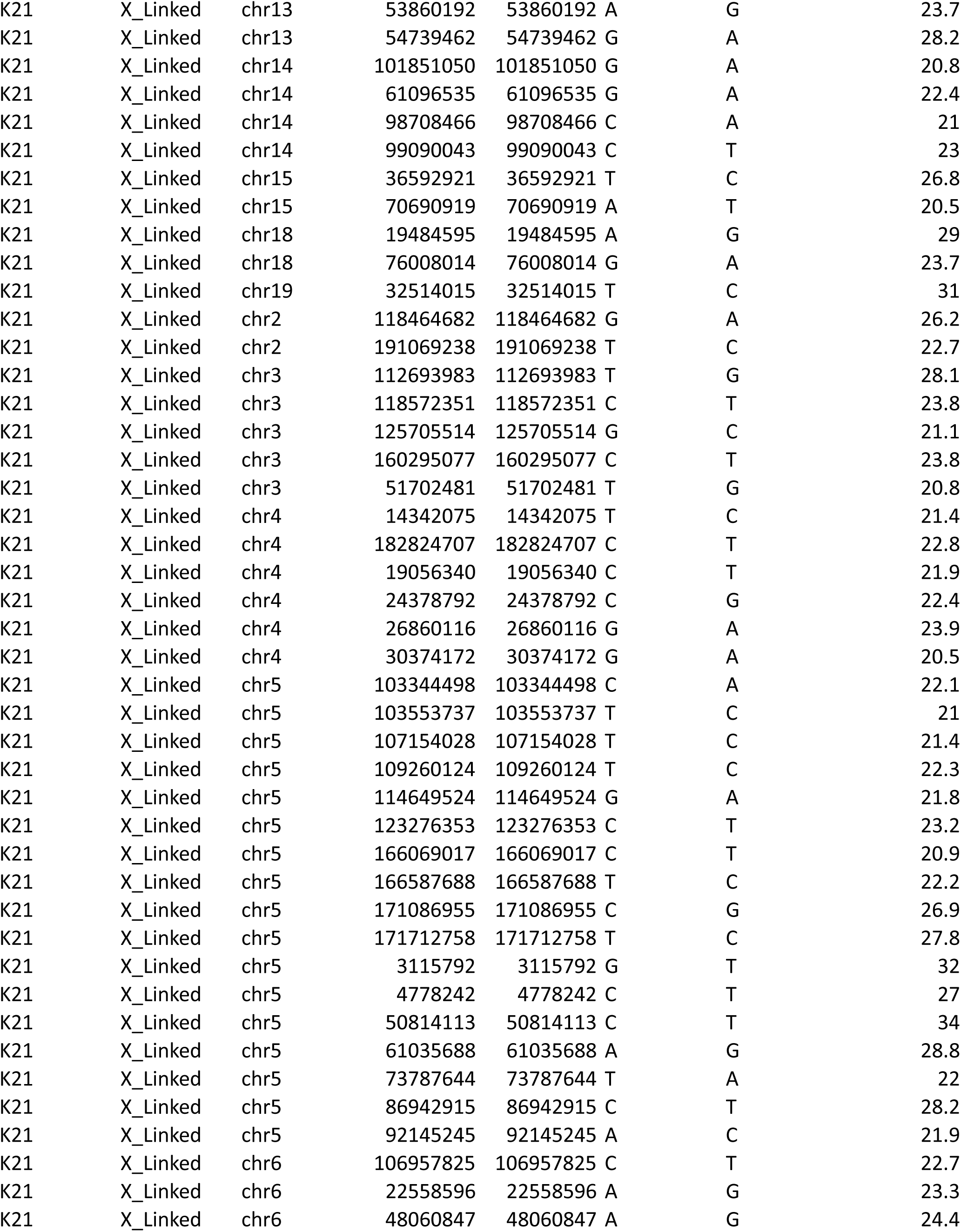
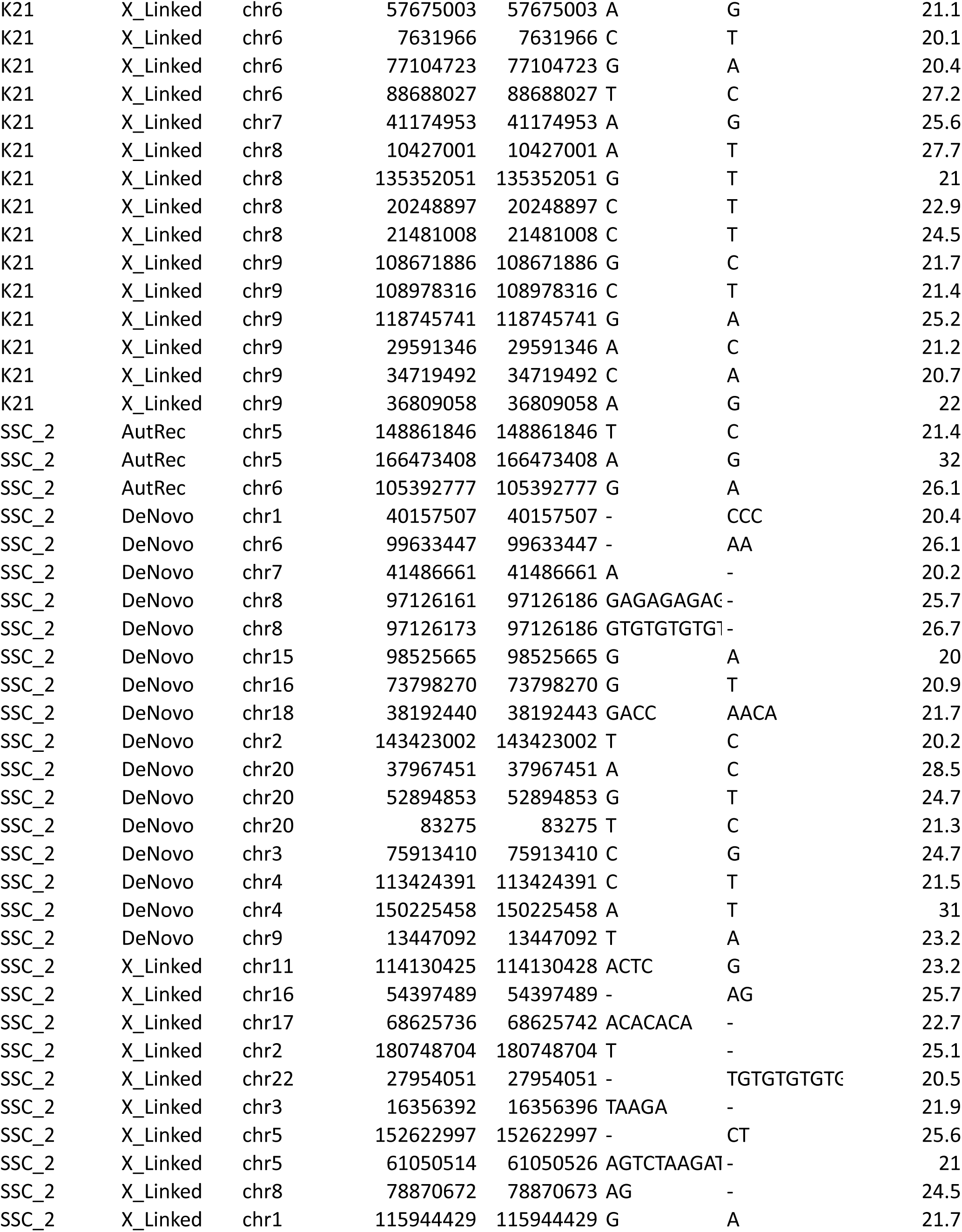
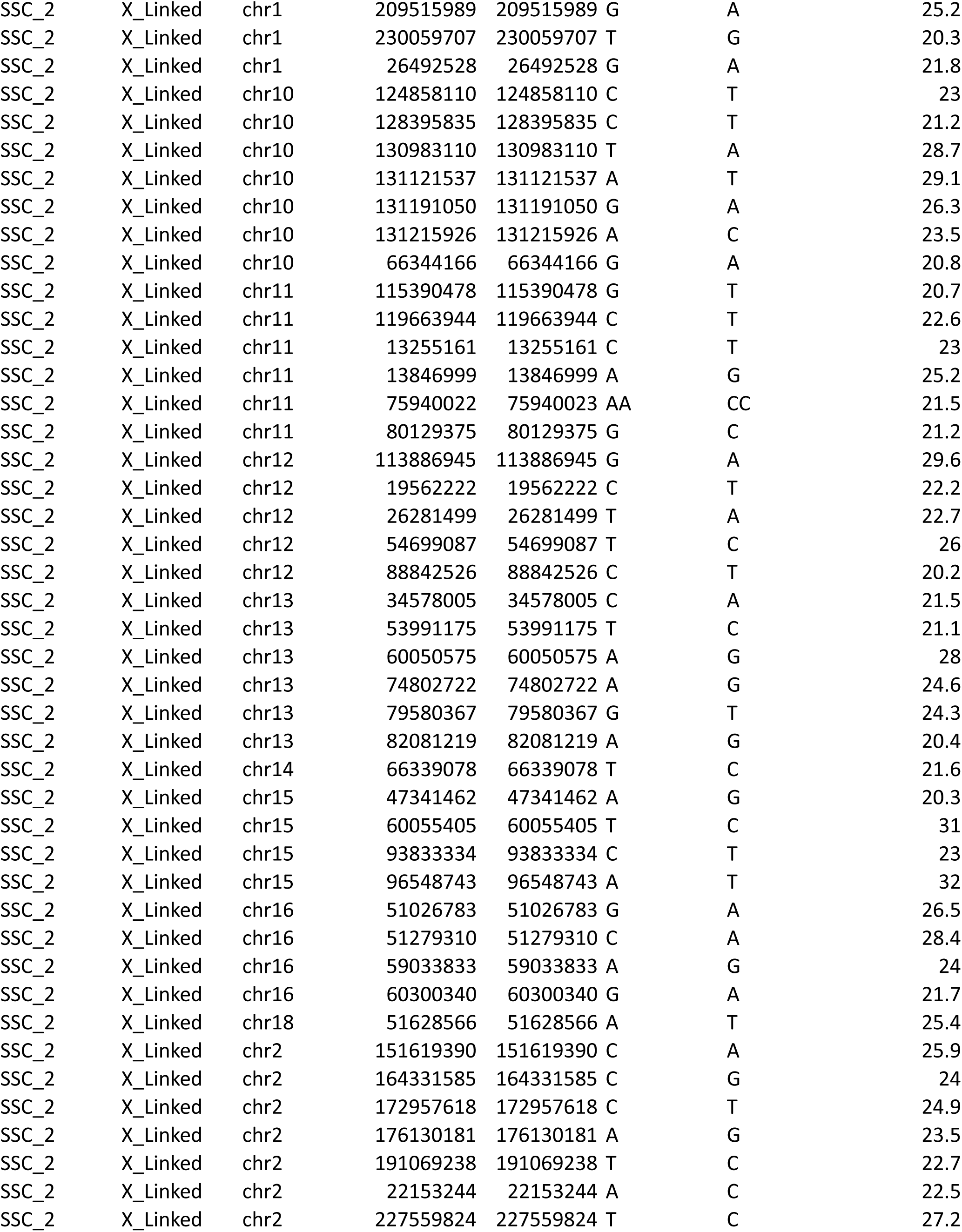
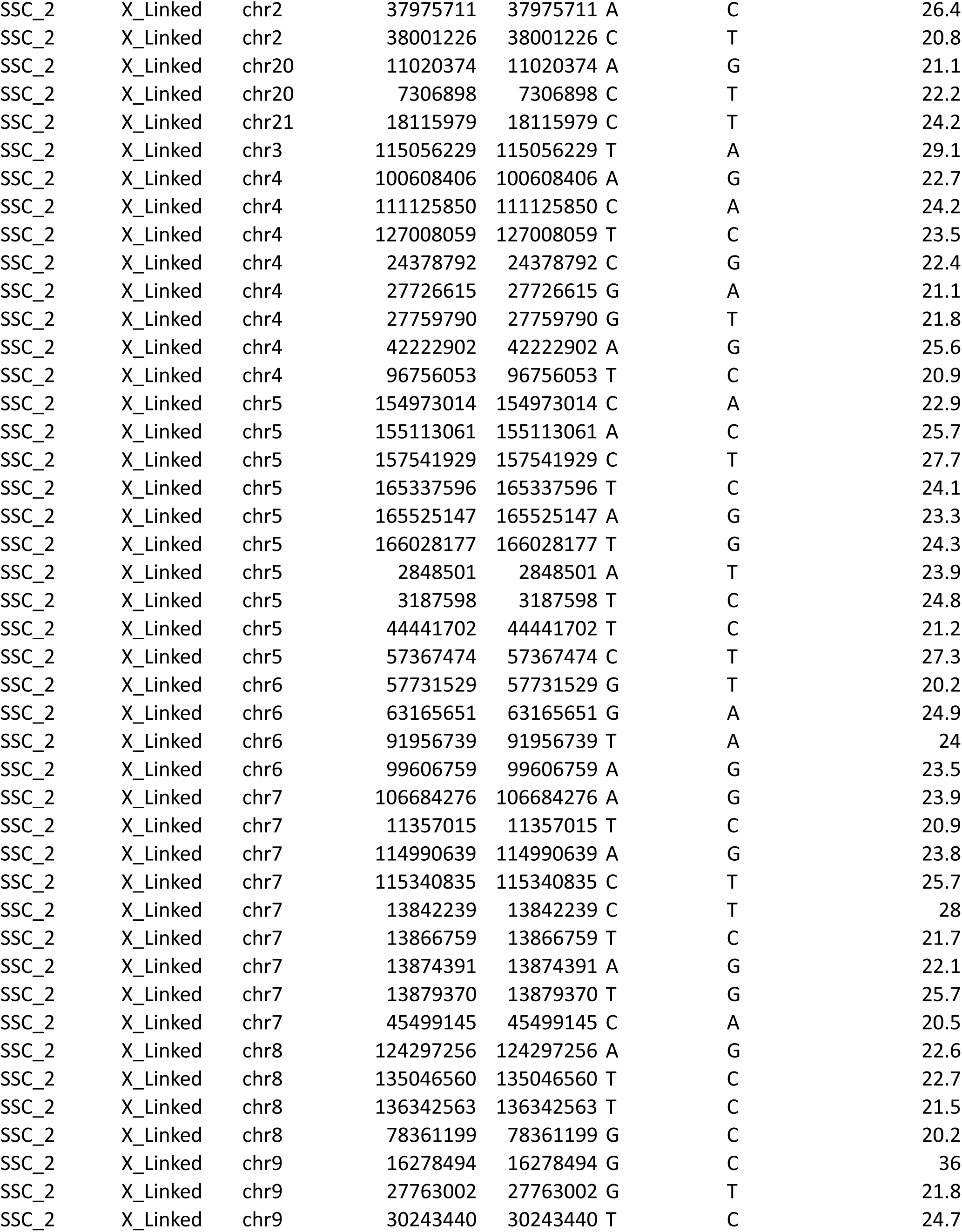
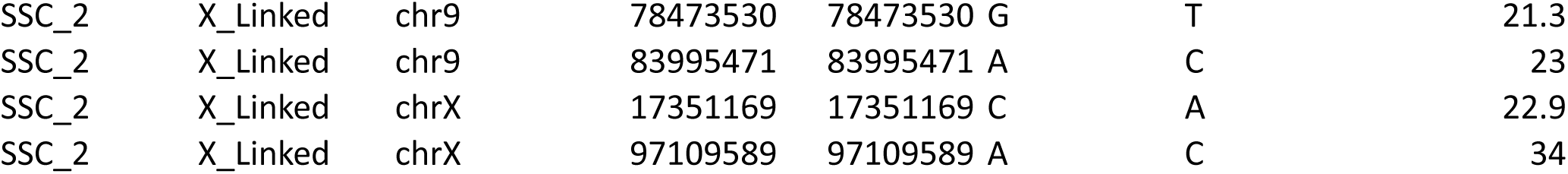

